# Soluble LAG-3 Identifies a Dynamic Early T Cell Activation Window in self-reactivity, Type 1 Diabetes, and Broader Immune Responses

**DOI:** 10.64898/2026.02.10.705171

**Authors:** Saptarshi Roy, Megan L. Proffer, Farooq Syed, Megan E. Smithmyer, Avinil Das Sharma, Abigail Twoy, Emily Soo-Hoo, Jyoti Rana, Jason M. Spaeth, Everett Meyer, Kent P Jensen, Cate Speake, William Hagopian, Jon D. Piganelli

**Author notes:** **Corresponding Authors**: **Jon D. Piganelli**, PhD, Department of Endocrinology, Indiana University School of Medicine 1210 Waterway, 4th floor, 4100F, Indianapolis, IN 46202., **Saptarshi Roy**, PhD, Department of Endocrinology, Indiana University School of Medicine 1210 Waterway, 4th floor, 41007, Indianapolis, IN 46202. Both authors equally contribute (Saptarshi Roy, Megan L. Proffer).

## Abstract

**Aims/hypothesis:** Type 1 diabetes is a complex autoimmune disorder in which autoreactive CD4⁺ and CD8⁺ T cells destroy pancreatic beta-cells, resulting in insulin deficiency and hyperglycemia. Although genetic susceptibility, particularly certain HLA alleles, contributes to disease risk, not all genetically predisposed individuals develop Type 1 diabetes. Screening first degree relatives (FDRs) for islet autoantibodies (GAD65, IAA, IA-2, ZnT8) helps detect autoimmune activity. However, these serum markers arise only after T-helper cell activation, limiting early intervention opportunities. Since protein antigen recognition by B cells requires T-helper cell assistance through linked recognition, T cell activation precedes B cell activation and autoantibody production. Activation of these T cells leads to shedding of the immune-regulatory (activation) surface protein LAG-3 (Lymphocyte Activation Gene-3 or CD223), generating its soluble form, sLAG-3, that is detectable in circulation. We hypothesized that sLAG-3 may serve as an early biomarker of autoimmune activity preceding islet autoantibody development in type 1 diabetes.

**Methods:** Plasma sLAG-3 levels were measured longitudinally in female diabetes-prone NOD mice and analyzed in relation to islet antigen-specific CD4⁺ T cell expansion and diabetes onset. To mechanistically link autoreactive T cell activation to sLAG-3 release. Naive autoreactive C6.6.9 TCR-transgenic (TCR-Tg) CD4⁺ T cells were adoptively transferred into NOD.SCID mice and longitudinal assessment for plasma sLAG-3, beta-cell antigen specific CD4⁺ T cell tetramer profiles, and circulating insulin (*Ins2*) mRNA to determine ongoing beta-cell stress. In parallel, sLAG-3 levels were analyzed from different human cohorts, including FDRs of individuals with type 1 diabetes, using cross-sectional and longitudinal approaches.

**Results:** In murine models, elevated sLAG-3 correlated with expansion of islet-specific CD4⁺ T cells that preceded hyperglycemia and diabetes onset. In the adoptive transfer model, early increases in sLAG-3 and circulating *Ins2* mRNA marked immune activation and emerging beta-cell stress prior to overt diabetes. In our human cohorts, sLAG-3 was detectable in autoantibody-negative and single-autoantibody-positive FDRs, with higher levels observed in progressors compared to non-progressors, and associated with high-risk HLA genotypes.

**Conclusions/interpretation:** These findings identify sLAG-3 as a candidate biomarker of early T cell activation in type 1 diabetes that may precede islet autoantibody development. Integration of sLAG-3 with antigen-specific T cell and beta-cell stress markers could improve early risk stratification and inform preventive strategies before substantial loss of beta-cell. Prospective longitudinal studies aligned to seroconversion are required to validate sLAG-3 as a surrogate marker of early disease activity.

**Research in context:** *What is already known about this subject?:*

- Before the clinical onset of hyperglycemia, type 1 diabetes is characterized by a prolonged preclinical phase in which autoreactive B and T cells mediate progressive beta-cell destruction.
- Current risk stratification strategies rely mainly on genetic susceptibility (genomic DNA) and the detection of islet autoantibodies in plasma/serum.
- Islet autoantibodies arise only after CD4⁺ T cell activation and therefore do not capture the earliest stages of immune dysregulation.
- Consequently, biomarkers that directly reflect early pathogenic T cell activity prior to, or independent of, seroconversion remain limited and insufficiently validated.

*What is the key question?:* Can plasma sLAG-3 levels, beta-cell antigen-specific CD4⁺ T cell tetramer expression, and circulating *Ins2* mRNA serve as very early biomarkers of autoimmune activity in type 1 diabetes and serve to better inform risk stratification, thereby informing preventive intervention strategies for the clinician?

*What are the new findings?:*

- sLAG-3 increases transiently during early antigen-specific CD4⁺ T cell activation stage, precedes hyperglycemia in mouse models, and is elevated in autoantibody-negative and single-autoantibody-positive first-degree relatives who later progress to type 1 diabetes.
- sLAG-3 was associated with beta-cell antigen-specific CD4⁺ T cell expansion, assessment of stress induced beta cell *Ins2* mRNA release and high-risk HLA genotypes, indicating early autoimmune activation rather than established disease.

*How might this impact clinical practice in the foreseeable future?:* These findings support sLAG-3 as a candidate early biomarker of T cell activation, before or at the earliest stages of islet autoantibody development in some at-risk individuals. Integration of plasma sLAG-3 with beta-cell antigen specific CD4⁺ T cell profiling and insulin mRNA measurements could complement current autoantibody-based screening, improve risk stratification, and enable earlier preventive interventions to preserve beta-cell function in patients at-risk for type 1 diabetes.

## Introduction

Type 1 diabetes is a complex, heterogeneous, and polygenic autoimmune disease characterized by CD4⁺, CD8⁺-T cell and B cell mediated destruction of pancreatic beta-cells, resulting in insulin deficiency and hyperglycemia [1–3]. Despite extensive research, the mechanisms driving this aberrant immune positive beta-cell interaction remain incompletely understood [4]. Type 1 diabetes affects 9.5 million people globally (2 million in USA). The global incidence is rising globally by 2-3% annually, outpacing the prevalence of high-risk HLA and other genetic alleles, indicating that environmental factors, including viral infections and oxidative stress, likely contribute to disease development [5–7]. First-degree relatives of type 1 diabetes patients, particularly those carrying HLA-DR3/-DQ2 and/or -DR4/-DQ8 HLA haplotypes, are at elevated genetic risk for type 1 diabetes. Patients and at-risk individual can be screened for islet autoantibodies in plasma/serum and HLA typing genomic DNA. Current biomarkers such as GAD65, insulin, IA-2 and ZnT8 autoantibodies are typically become detectable only after substantial beta-cell loss, indicating a clinically silent window of early immune dysregulation that remains undetected [8, 9]. Thus, there is a crucial and yet unmet need to validate this silent window for biomarkers capable of identifying type 1 diabetes at the earliest pre-seroconversion stages of autoimmunity.

The FDA approval of teplizumab, an anti-CD3 lymphodepleting immunomodulatory therapy, represents a major advance in our field that, demonstraties that disease onset can be delayed when treatment is initiated post-autoantibody seroconversion [10]. Nevertheless, full disease prevention remains challenging, as intervention at this stage occurs after significant beta-cell injury [11]. Since seroconversion occurs only after immune tolerance is breached and self-reactive CD4⁺ T cells are activated, earlier biomarkers that detect pathogenic immune activity and distinguish progressors from non-progressors are urgently needed to inform prevention strategies [11–13]. Autoantibodies primarily reflect loss of immune tolerance but arise only after CD4⁺ T cells provide help for B cell activation and antibody production [10, 11]. Early antigen-specific CD4⁺ T cells deliver essential co-stimulatory signals and cytokines that drive B cell proliferation, isotype class switching, and differentiation into plasma cells that produce high-affinity autoantibodies against beta-cell antigens [14]. While both genetic and environmental factors likely contribute to CD4⁺ T cell activation, the initiating triggers remain unclear [15, 16].

During T cell activation, a cellular metalloprotease (ADAMs-17) enzymatically cleaves the lymphocyte activation gene-3 protein from the cell surface, releasing sLAG-3 into circulation as a T cell activation marker [12, 14, 15]. LAG-3 is a co-inhibitory receptor expressed on regulatory T cells that is structurally similar to CD4 [17]. Increase sLAG-3 levels, previously associated with strong immune responses in cancer, and Graves’ Disease, appears to predict emerging autoimmunity in type 1 diabetes prior to autoantibody detection [16, 18, 19]. Concurrently, in beta cells proinflammatory stress mediate release of *insulin* mRNA into circulation, representing another promising early marker of beta-cell dysfunction [20–22]. However, the lack of early biomarkers that can be used to identify at-risk individuals before seroconversion, highlights the need for enhanced assays capable of detecting both sLAG-3 and *insulin* mRNA to monitor early immune activity and beta-cell health.

To evaluate the feasibility of these proposed biomarkers panel, we conducted a preliminary study integrating plasma sLAG-3 quantification via monoclonal antibody-based ELISA assay with beta-cell antigen-specific CD4⁺ T cell tetramer profiling. This combinatorial assay was applied to longitudinal plasma samples from female diabetic-prone NOD mice, a human cohort from the Diabetes Evaluation in Washington (DEW) study, and cross-sectional autoantibody-positive samples from the Stanford Diabetes Biobank. In these preclinical and clinical samples, we observed early increases in plasma sLAG-3 coincided with rising frequencies of beta-cell antigen-specific CD4⁺ T cells. Our results accurately predicting impending diabetes onset and supporting the potential utility of this biomarker combination. We further examined early immune activation using a longitudinal adoptive-transfer model of autoimmune diabetes in which naïve islet-reactive CD4⁺ T cells from NOD.C6.6.9 TCR-transgenic mice were transferred into immunodeficient NOD.SCID recipients. This reductionist system enables real-time monitoring of autoreactive T cell antigen engagement, activation, expansion, antigen engagement, and effector differentiation. We assessed plasma sLAG-3 concentration, CD4⁺ T cell tetramer positivity, and circulating *insulin* mRNA levels as biomarkers of beta-cell health and disease progression [20]. Notably, the early rise in circulating *Ins2* mRNA in whole blood sapless preceded hyperglycemia and coincide with increases in immune cell infiltration into the pancreatic islets, suggesting that immune-driven β-cell dysfunction disrupts insulin protein synthesis and secretion far earlier than previously recognized [23–25]. These findings support the use of metabolic markers to detect early beta cell vulnerability, that could complement immune-activation biomarkers of immune activation and uncover targets to preserve insulin secretion. Additionally, human studies further confirm an increase in sLAG-3 in plasma of single autoantibody-positive first-degree relatives of individuals with type 1 diabetes compared to individuals who are autoantibody-negativity, reinforcing its potential as an early biomarker. Combined monitoring of sLAG-3 and CD4⁺ T cell tetramer expression, along with circulating *Ins2* mRNA, may enable earlier type 1 diabetes detection and inform type 1 diabetes intervention trial before substantial beta-cell loss and clinical onset [20, 24].

## Methods

### Animal Models and Diabetes Assessment

NOD/ShiLtJ (Strain 001976), NOD.Cg-Prkdc^scid/J^ (Strain 001303), and C57BL/6 (Strain 000664) male and female mice were obtained from Jackson Laboratory. Beginning at 4 weeks of age, NOD/ShiLtJ mice were routinely monitored for glucosuria in 3-day intervals during adoptive transfer studies. Blood glucose was measured in mice positive for glucosuria, and overt diabetes was defined as two to three consecutive readings >300 mg/dL. Blood glucose was measured by puncturing the last 1 mm of the tail and dispensing one drop of blood on a Contour next® disposable test strip and glucometer. All mice were bred and maintained under specific pathogen-free conditions at Indiana University’s BRTC, managed by LARC. Animal experiments were conducted in accordance with Animal Research: Reporting of In Vivo Experiments (ARRIVE) guidelines under protocols approved by the Indiana University (IU) School of Medicine Animal Use Committee.

### Research participant’s samples

Plasma from healthy controls, type 1 diabetes participants, and FDRs were provided by Children’s Hospital of Pittsburgh, Stanford University School of Medicine (IRB Protocol #35453), University of Florida, and University of Queensland. Samples were stored at −80°C until analyzed. Participants with type 1 diabetes ranged in age at onset (1 to 16 yrs old) and duration of disease (1 to 6 years post-onset). FDRs were tested for auto-antibodies to GAD65, IAA, and IA2 by standard radioimmunoassay. Plasma samples were collected from two cohorts of healthy controls, younger participants aged 25-35 and older participants aged 55-65 years old who were recruited into the Sound Life Project for measuring the sLAG-3 level from research participant before and after covid vaccination. Samples were collected with approval from Benaroya Research Institute, Seattle, WA review board (IRB19-045) and the Stanford University School of Medicine institutional review board (IRB 35453). All participants provided written informed consent. Prospective cohort longitudinal studies for measuring the sLAG-3 from Diabetes Evaluation in Washington Study (DEWIT) enrolled infants from the general population (n=39) carrying increased-risk HLA-DR-DQ genotypes identified by newborn screening, as well as first-degree relatives of individuals (n=13) with or without type 1 diabetes regerdless of HLA genotype. All participants underwent HLA screening, though eligibility criteria varied across sites. Follow-up began between 2 months and 21.6 years of age and continued at 3 to 36 month intervals for up to 26 years, with type 1 diabetes diagnosed according to American Diabetes Association criteria. All protocols were approved by local institutional review boards, and de-identified data were submitted to IBM Research. The harmonized cohort comprised median 10 visits, 8.7 years follow-up; 17% reported a family history of type 1 diabetes. For key analyses, it was examined the infant-toddler subcohort, restricted to subjects tested for insulin, IA-2, and GAD autoantibodies by age 2.5 years. The infant-toddler cohort included median 12 visits, 10.4 years follow-up [26].

### Estimating soluble LAG-3 levels from NOD and human plasma

NOD/ShiLtJ and NOD.Cg-Prkdc^scid/J^ mice from different experimental groups were retro-orbitally bled for plasma every other week from 4 to 16 weeks of age. Plasma was isolated by centrifugation at 3000 × RPM for 10 minutes at 4°C. Mouse sLAG-3 was measured by sandwich ELISA using 96-well high-affinity plates (Corning #3361) coated overnight at 4°C with 5 µg/mL monoclonal anti-LAG-3 (C9B7W, InVivoMab #BE0174). LAG-3 standard was purchased from R&D (Catalog: AF3328) antibody pairs for mouse compatible were purchased from Jackson Immuno Research, performed according to manufacturer’s protocol. Human plasma from three institutions was analyzed using Human LAG-3 DuoSet ELISA (R&D #DY2319B). All samples were run in triplicate, read at OD450 nm on a SpectraMax-iD5, and analyzed with SoftMax-Pro 7.0.2.

### Plasma sLAG-3 evaluation from DEWIT cohort study and Stanford Biobank

Human plasma samples were analyzed for sLAG-3 levels by the MSD U-PLEX platform using sandwich immunoassays for protein targets and a competitive assay for neopterin, with electrochemiluminescence detection. MULTI-ARRAY® 96-well plates contained sLAG-3 capture antibodies, Sulfo-Tag detection antibodies, calibration standards, and diluents. The Indiana University Translational Core processed 50 μL of each sample in triplicate. Mean concentrations were reported, with values below the limit of detection (LOD) assigned the LOD. Plate-specific control coefficients of variation (CVs) were calculated for sLAG-3.

### Estimating sLAG-3 from covid vacationed research participants

sLAG-3 level from covid vacationed research participants plasma was measured using the Olink proximity extension assay on the Olink Explore 1536 platform’s oncology panel [27]. Samples were randomized across plates to balance age and sex, with longitudinal samples from the same participant run on the same plate. Bridging controls were included in each batch for cross-batch normalization. Protein expression values were first normalized within wells using internal extension controls (IgG antibodies with matched oligos), then standardized across plates using triplicate inter-plate pooled serum controls. Data were subsequently intensified across all samples, and final normalized relative protein quantities were reported as NPX values.

### In vivo viral infection incidence study

Eight-week-old C57BL/6 mice were intraperitoneally infected with 100 PFU Coxsackievirus B (CVB3 #ATCC® VR-30™) in HBSS under specific pathogen-free conditions. Blood was collected via retro-orbital bleeding at baseline and on days 3, 6, 9, 12, 16, and 21 post-infections. Plasma was isolated by centrifugation and stored at -80°C. sLAG-3 levels were quantified using standard ELISA, while intracellular IFN-γ in CD4⁺ and CD8⁺ T cells was measured by flow cytometry to assess immune activation during infection.

### Adoptive transfer

Spleens from 8-week-old female NOD congenic C6.6.9 TCR-Tg mice were harvested sterilely, mechanically homogenized, passed through a 40 µm strainer, washed, and resuspended in cold DMEM with 10% FBS. Lymphocytes were isolated via Ficoll-Paque density gradient, washed twice with PBS, and 5×10⁶ cells were retro-orbitally transferred into 8-week-old female NOD.Cg-Prkdc^scid/J^ recipients. Post-transfer, mice were monitored for diabetes progression by blood glucose and bled every three days to measure plasma sLAG-3, activation markers and co-inhibitory receptor (CD69, CD44, LAG-3), and CD4⁺ T cell reactivity to beta-cell antigens (ChgA, InsB9-23, IAPP) using tetramers. Mice with >300 mg/dL glucose on two readings were euthanized for spleen and pancreas for further analyses.

### Staining and flow cytometry analysis

To assess T cell mediated immune activation in autoimmune and viral contexts, lymphocytes were analyzed by flow cytometry. NOD/ShiLtJ mice were longitudinally bled during diabetes progression, and NOD.SCID mice were sampled after adoptive transfer. For viral studies, CD4⁺IFN-γ⁺ and CD8⁺IFN-γ⁺ T cells were analyzed from C57BL/6 blood. Lymphocytes were isolated, RBCs lysed, and cells resuspended in FACS buffer. Cells were stained with eFluor-780 viability dye and surface markers including CD45-FITC, CD4-PerCP-Cy5.5, CD8-APC, CD69-PE/Cy7, CD25-PE, LAG-3-BV650, CD71-APC, CD98-PE/Cy7, and Glut-1-PE. For beta-cell antigen specific, APC-ChgA, BV421-InsB9-23, and PE-IAPP tetramers were used. Intracellular IFN-γ-PE staining was performed using the Fixation & Permeabilization Buffer Kit protocol (eBiosciences FOXP3 Fix/Perm kit, Cat #00-5523-00). Data were analyzed using FlowJo-v10.

### Cytokine measurement by ELISA

Plasma from CVB-infected C57BL/6 mice was collected at multiple time points and analyzed for IFN-γ by ELISA. High-affinity 96-well plates were coated overnight at 4°C with 25 ng/mL monoclonal rat anti-murine IFN-γ in carbonate buffer (pH 9.6). Mouse IFN-γ ELISAs were performed using BD antibody pairs following the manufacturer’s protocol. Mean OD450 values were recorded on a SpectraMax-iD5 microplatereader, analyzed using SoftMax Pro-7.0.2.

### Histology and Immunofluorescence

Pancreata from control, diabetic, and post-adoptive transfer NOD.SCID mice (days 12 and 15) adoptively transferred NOD.SCID mice were harvested at diabetes onset, fixed in 4% paraformaldehyde for 4 hours at 4°C, washed with distilled water, and processed for paraffin embedding, and stained with hematoxylin and eosin to assess tissue architecture and immune infiltration following standard protocol. For immunofluorescence, pancreata cryoprotect overnight at 20% sucrose in PBS. Tissues were embedded in OCT, frozen, and stored at -80°C. Cryosections (6 µm) were prepared using a cryostat. Sections were blocked with 5% normal donkey serum and 1% BSA, incubated overnight at 4°C with monoclonal CD3, insulin, F4/80, and B220 primary antibody, followed by fluorophore-conjugated secondary antibodies. Similarly, APC-ChgA, BV421-InsB9-23, and PE-IAPP tetramers antibody are also used. Nuclei were counterstained with DAPI. Images were acquired on a Zeiss LSM-800 and analyzed using ImageJ.

### RNA isolation and ddPCR

Total RNA was isolated from the blood of adoptively transferred NOD.ScidSCID mice using RNAprotect Animal Blood Tubes (cat. #76544) and the RNeasy Protect Animal Blood Kit (cat. #73224) per manufacturer instructions. RNA concentration was measured with a NanoDrop-2000 (Thermo Fisher Scientific). cDNA was synthesized using M-MLV Reverse Transcriptase (Invitrogen) according to the manufacturer’s protocol. Droplet digital PCR (ddPCR) was performed to quantify *Ins2* mRNA. Briefly, 2 μL of cDNA was combined with 10 μL ddPCR Supermix for Probes (1863024) and 1 μL TaqMan primer (Cat #4427975) that target *Ins2* transcript and the volume was adjusted to 20 μL with RNase/DNase-free water. Synthetic *Ins2* cDNA was used as a positive control and RNAse free water was used negative controls. Droplets were generated with QX200 AutoDG, PCR on a C1000 Thermal Cycler, and read with QX200 droplet reader. Data (copies/μL) were analyzed using QuantaSoft software.

### Statistical Analysis

Data analysis was performed using GraphPad Prism10. Pairwise comparisons were performed using unpaired or paired two-tailed Student’s t-tests. Interactions between two groups with two variables (e.g., time or treatment, pre-post-vaccination) were determined using a one-way ANOVA followed by an appropriate post hoc test. Sample sizes (n), means, error bars, and statistical methods are described in figure legends. A p-value <0.05 indicates statistical significance.

## Results

### Plasma sLAG-3-levels track spontaneous diabetes onset in NOD mice

LAG-3 (CD223) is a co-inhibitory receptor expressed by CD4+CD25+CD127lo regulatory T (Treg) cells and other regulatory immune cell subsets [28, 29]. We previously proposed that protein enzyme catalytic liberation of LAG-3 from the surface of CD4 Tregs (solubilized or sLAG-3) is a key biomarker, indicative of loss of tolerance to pancreatic islet beta cell antigens in autoimmune-prone NOD mice [30]. To determine whether soluble LAG-3 In plasma/serum reflects early autoimmune activation, we followed female NOD mice longitudinally from 4-16 weeks of age, measuring plasma sLAG-3 and blood glucose biweekly (Fig.1a). Plasma sLAG-3 was low at 4 weeks of age, increased significantly during 6 to12 weeks, reflecting early T cell activation and then declined to near-baseline levels by 16 weeks of age (Fig. 1b). This window of elevated sLAG-3 coincided with the prediabetic period when insulitis is known to expand in NOD mice [31]. In contrast, blood glucose remained normal through 12 weeks and began to rise around 16 weeks of age as overt, clinical diabetes appeared (Fig. 1c). Importantly, a significant positive correlation was observed between peak sLAG-3 levels and the timing of diabetes onset in the female NOD cohort (Fig. 1d), indicating that higher early sLAG-3 levels are associated with greater likelihood of progression to hyperglycemia in this model.

**Fig. 1.**
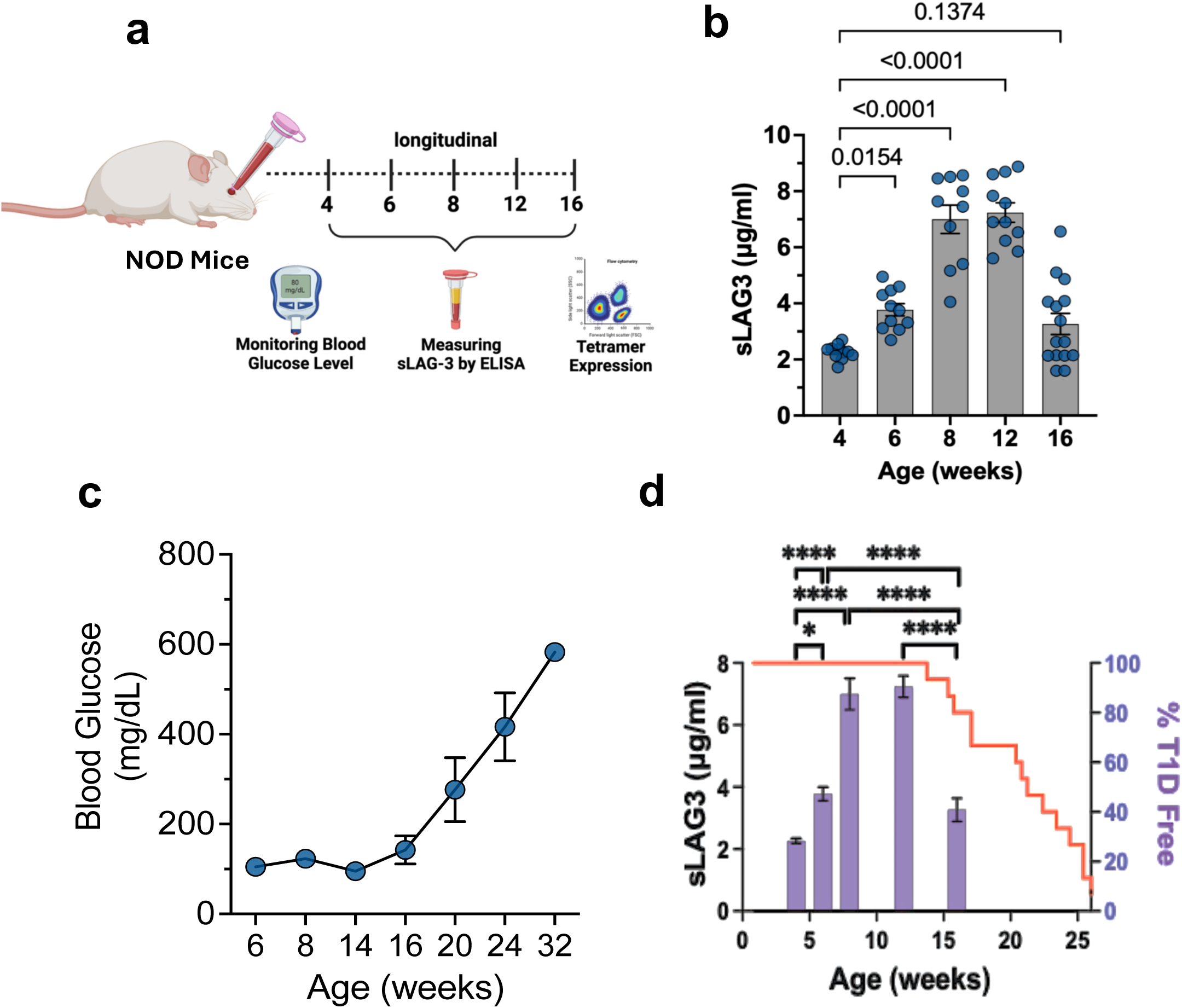
Plasma sLAG-3 correlates with disease incidence in NOD mice. (**a**) Experimental design for longitudinally monitoring blood glucose and plasma sLAG-3 level from female NOD mice. (**b**) Plasma sLAG-3 levels in NOD mice at different ages. (**c**) Longitudinal monitoring of blood glucose levels. (**d**) Correlation between plasma sLAG-3 levels and diabetes incidence in NOD mice. Data are presented as mean ± SEM (n=10-12 per group). Statistical significance was determined using one-way ANOVA followed by Bonferroni post-hoc testing; *\*p<0.05*.

### Beta cell antigen-specific CD4⁺ T cells expand in parallel with sLAG-3 during spontaneous disease

We next examined whether beta-cell antigen specific CD4⁺ T cells show similar temporal dynamics during spontaneous disease in female NOD mice from the same longitudinal cohort (Fig. 1a). Using MHC class II tetramers for the pancreatic islet beta cell antigens InsB923, ChgA, and IAPP, we observed-a significant increase in both the frequency and absolute number of tetramer positive CD4⁺ T cells between 6 and 12 weeks of age (Fig. 2a, b). Mean fluorescence intensity (MFI) of tetramer staining also increased over this interval (Fig. 2c), consistent with either higher TCR occupancy or enrichment of higher-affinity clones. Among the three specificities, InsB9-23⁺ CD4⁺ T cells were the most abundant at 6 weeks and increased further by 12 weeks of age. In contrast, ChgA and IAPP-specific cells started at a lower frequency and expanded later time points, suggesting a progressive broadening of beta cell antigen recognition within the CD4⁺ compartment. However, consistent with Fife et al. findings, reveal that in spontaneous diabetes, multiple autoreactive T cell populations infiltrate the islets, including ChgA-specific BDC2.5, and IAPP-reactive BCD6.9 T cells. Among these, BDC2.5 T cells are the most pathogenic, likely owing to the robust production of IFN-g and TNF-a [32], whereas insulin-reactive T cells, despite their abundance, exhibit limited immunopathogenicity [33]. The rise in tetramer-positive cells coincided with the period of elevated plasma sLAG-3 and preceded the clinical onset of hyperglycemia (Fig. 1b, c, and 2d). Linear regression analysis confirmed a significant positive association between plasma sLAG-3 levels and the frequency of beta-cell antigen–specific CD4⁺ T cells in our NOD murine studies (Fig. 2e), linking systemic sLAG-3 shedding to antigen-specific T cell activation and loss of tolerance to beta cells and their protein antigen during early autoimmune progression.

**Fig. 2.**
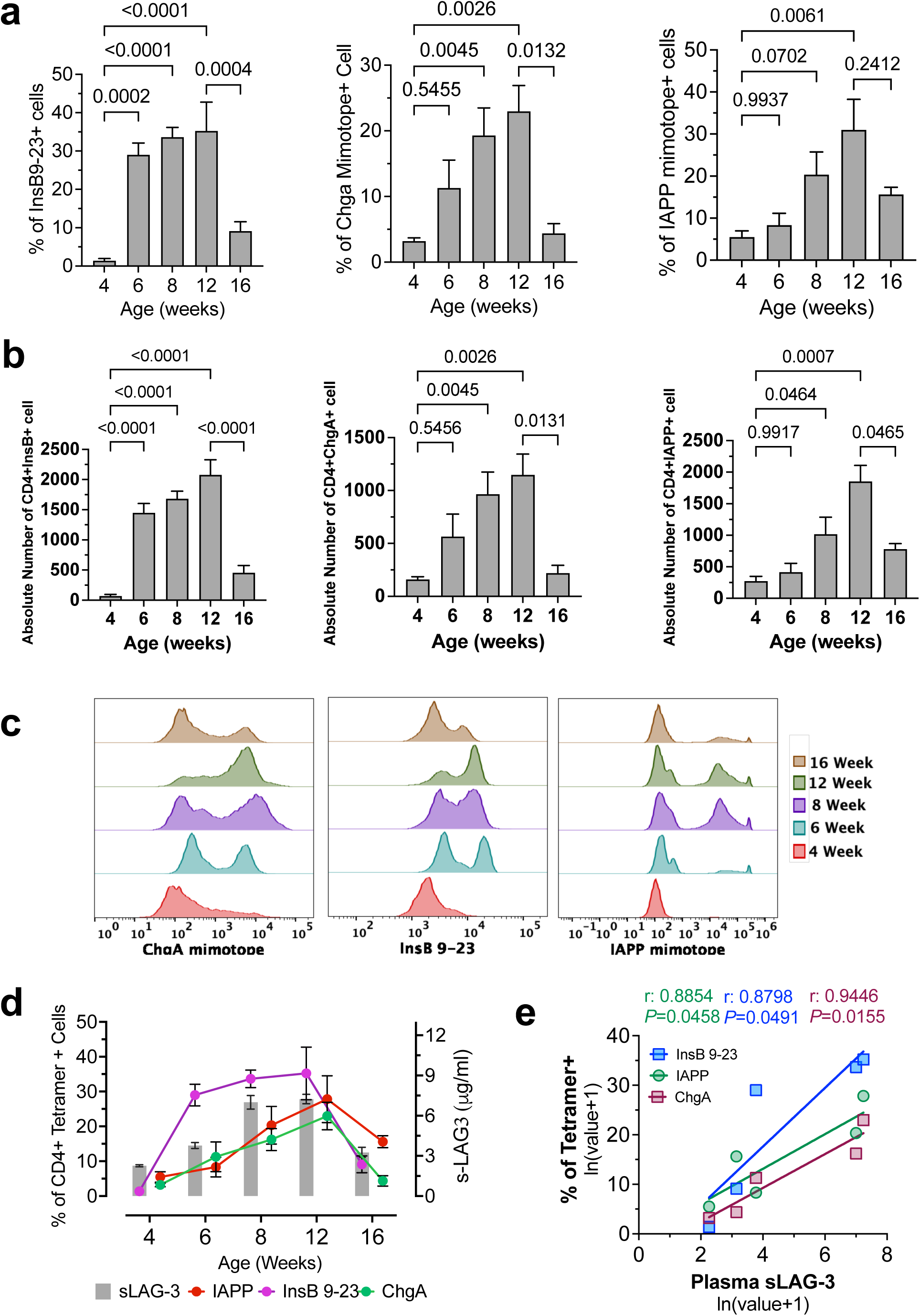
Beta-cell antigen-specific tetramer expression in CD4⁺ T cells corelate with disease incidence in NOD mice. (**a**) Percentage of CD4⁺ tetramer positivity T cells for three islet antigens (InsB9-23, ChgA, and IAPP) (**b**) Absolute numbers of tetramer-positive CD4⁺ T cells. (**c**) Overlay of the MFI histograms by flow cytometry analysis of tetramer expression in CD4⁺ cells (**d**) Overlay representation of plasma sLAG-3 and CD4⁺ tetramer expression. (**e**) Linear regression analysis of plasma sLAG-3 levels and islet antigen-specific tetramer positive CD4⁺ T cell frequencies in CD4⁺ T cells in NOD mice during diabetic progression. Data are presented as mean ± SEM (n=10-12 in the study cohort) Statistical significance was determined using one-way ANOVA with Bonferroni post-hoc testing; *\*p<0.05*.

### Adoptive transfer of C6.6.9 CD4⁺ T cells reveals early sLAG-3 induction and beta-cell stress

To further study the relationship between beta-cell injury, beta-cell antigen–specific CD4⁺ T cells activation/expansion and release of sLAG-3 release and beta-cell injury, we adoptively transferred splenocytes from NOD.C6.6.9 TCR transgenic mice into NOD.SCID recipients (Fig. 3a). Donor mice express the BDC-6.9 TCR but lack the cognate IAPP-derived hybrid peptide, so their CD4⁺ T cells remain naïve prior to transfer. After adoptive transfer, recipient NOD.SCID mice undergo a prediabetic phase followed by a rapid metabolic -shift with in, blood glucose began to rise by approximately 7 weeks after transfer and exceeded 400 mg/dL by week 10 (Fig. 3b, c and ESM Fig. 2b). Plasma sLAG-3 levels increased early following adoptive transfer, peaking between days 12-15 after adoptive transfer and then declining as diabetes developed (Fig. 3d; ESM Fig. 2a). In line with the data from sLAG-3 secretion, circulating levels of *Ins2* mRNA showed a similar early increase (Fig. 3e), indicating ongoing beta-cell stress during the same interval. *Ins2* mRNA displayed a modest positive association with sLAG-3 (Fig. 3f), but neither marker correlated positively with blood glucose, consistent with the idea that substantial immune and beta-cell stress precede overt hyperglycemia (Fig. 3g). However, sLAG-3 showed an inverse correlation with blood glucose when later time points were included (Fig. 3h), indicating high sLAG-3 in circulation early during autoreactive T cell activation followed by a decline as beta-cell mass is lost. Flow cytometry revealed dynamic redistribution of donor C6.6.9 CD4⁺ T cells-over time, with expansion in the periphery followed by accumulation in the pancreas and secondary lymphoid organs (Fig. 3i). Histopathological analysis of pancreata from diabetic recipients showed marked signs of immune cell infiltration and insulitis, with extensive islet destruction (Fig. 3j). Together, these data confirm that beta cells antigen specific CD4⁺ T cells are sufficient to drive diabetes in this model and that early activation is accompanied by a transient rise in plasma sLAG-3 and beta-cell *Ins2* mRNA.

**Fig. 3.**
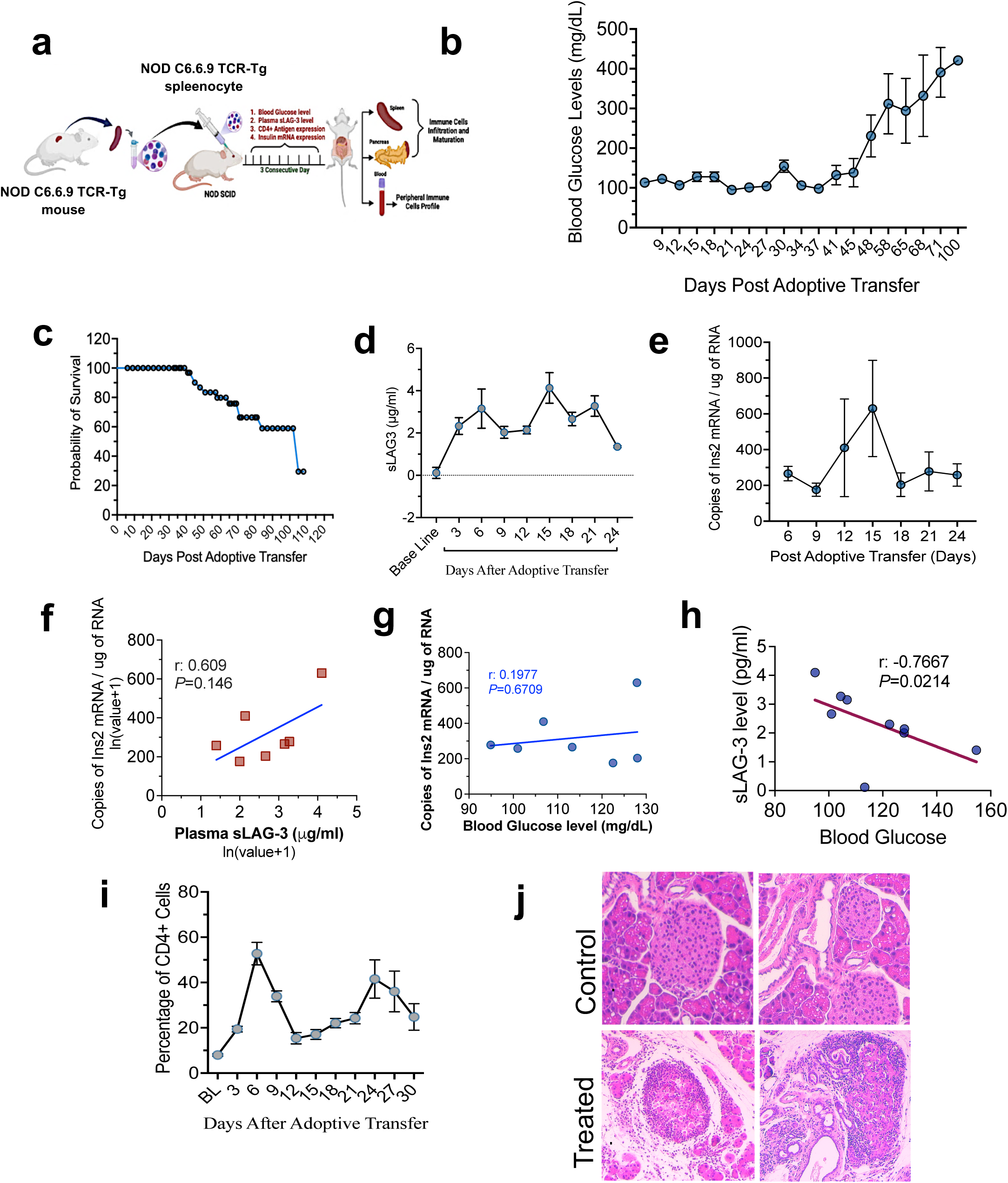
Antigen-specific CD4⁺ T cell activation and immune modulation during diabetes progression following adoptive transfer and immune modulation. (**a**) Schematic overview of the experimental design for adoptive transfer and tracking activation of naive C6.6.9 TCR-Tg CD4⁺ T cells in NOD.Scid recipient mice. (**b**) Longitudinal monitoring of blood glucose levels post-transfer. (**c**) Kaplan-Meier survival analysis of diabetes onset following T cell transfer. (**d**) Plasma sLAG-3 levels. (**e**) Plasma *Insulin2 (Ins2)* mRNA levels measured at defined time points post-transfer. (**f-h**) Linear regression analyses relating plasma sLAG-3 levels, plasma Ins2-mRNA, and blood glucose levels during diabetes progression. (**i**) Frequency of circulating C6.6.9 TCR-Tg CD4⁺ T cells in peripheral blood of recipient mice over time following adoptive transfer. (**j**) Representative histological images showing progressive insulitis and islet destruction in pancreatic tissue from NOD.Scid recipient mice. Data are presented as mean ± SEM (n=10).

### Longitudinal assessment of antigen specific CD4⁺ T cell activation and beta-cell antigen tetramer profiles in adoptively transferred mice

We next characterized the activation state and antigen specificity of CD4⁺ T cells in peripheral blood of adoptively transferred NOD.SCID mice every three days. Shortly after transfer, CD4⁺ T cells exhibited a pronounced activation surge defined by increased CD25⁺ and CD69⁺ frequencies, peaking between days 3-9 following adoptive transfer (Fig. 4a; ESM Fig. 3a). Concomitantly, the frequency of ChgA⁺, InsB9-23⁺, and IAPP⁺ tetramer-positive CD4⁺ T cells transiently rose between 9 and 15-days post-transfer and then declined as cells redistributed into tissues (Fig. 4b; ESM Fig. 3b). LAG-3 expression on CD4⁺ T cells also increased early and then decreased among adoptively transferred CD4 T cells, consistent with the pattern of soluble LAG-3 detected in plasma. Therefore, to address the mechanism underlying the emergence of these newly detected antigen-specific T cell populations, we performed immunofluorescence staining of pancreata at days 12 and 15 post adoptive transfer. We found that tetramer-positive cells, particularly -ChgA⁺, were more abundant at day 12 than day 15 (Fig. 4c, d), mirroring the transient early peak observed in blood. At diabetes onset, analysis pancreatic tissue revealed that CD69 expression was significantly increased on pancreatic CD4⁺ T cells compared with blood. In contrast, CD25 expression was relatively higher in circulating CD4+ T cells (Fig. 4e; ESM Fig. 3c). Tetramer positive CD4⁺ T cells specific for ChgA, InsB9-23, and IAPP were most enriched in the pancreas compared with spleen and blood (Fig. 4f; ESM Fig. 3d), indicating preferential homing and/or retention of autoreactive CD4⁺ T cells at the site of beta-cell destruction. Consistent with findings by Bettini and Vignali et al., T cells that are not specific for islet beta-cell antigens do not persist within the pancreatic islets and exit once cognate antigen recognition fails to occur [34].

**Fig. 4.**
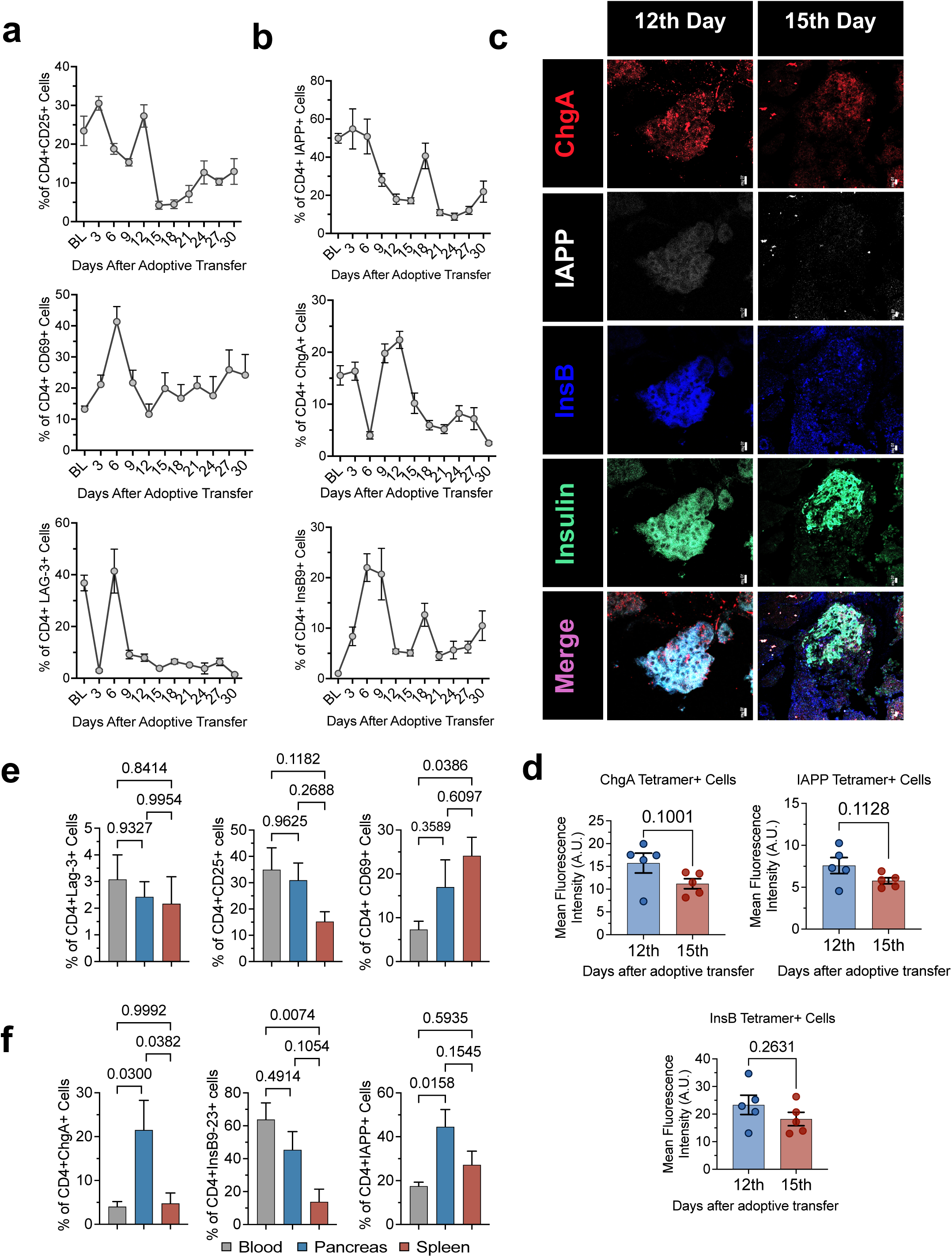
Longitudinal phenotypic characterization of T cell activation markers in recipients following adoptive transfer. (a) Frequency of T-cell activation markers (CD25, CD69, and surface LAG-3). (b) Frequency of β-cell antigen-specific tetramer-positive CD4⁺ T cells in peripheral blood over time. (c) Immunofluorescence staining of pancreatic sections on days 12 and 15 post-transfer showing infiltration of ChgA, InsB9-23, and IAPP-reactive T cells. (d) Quantification of immunofluorescence MFI for ChgA⁺, InsB9-23⁺, IAPP⁺ cells from pancreatic sections. (e, f) Percentage of different activation markers (CD25, CD69, LAG-3) and β-cell antigen-specific tetramers (ChgA, InsB9-23, IAPP) expression on T cells that had migrated or infiltrated into blood, pancreases and spleen at the time of diabetes. All those values are represented as mean ± SEM (n=4-10). P <0.05 is significant and P>0.05 not significant. Statistics were done by one-way ANOVA with multiple comparisons, followed by the Bonferroni post hoc test.

### Beta-cell antigen driven metabolic reprogramming and immune infiltration in the pancreas

To assess metabolic changes accompanying antigen driven T cell activation, we examine the immune infiltration in pancreas and quantified nutrient transporter expression on CD4⁺ T cells isolated from blood, spleen, and pancreas at diabetes onset in adoptively transferred NOD.SCID mice (Fig. 5a). Immunofluorescence confirmed dense infiltrates of CD3⁺ T cells along with B220⁺ B cells and F4/80⁺ macrophages in diabetic pancreata compared with controls, together with loss of insulin positive beta-cells (Fig. 5b), suggests that autoreactive T cells maybe directly and indirectly by activating macrophage the local milieu to support B cell help and drive metabolic programs sustaining effector function [35–37]. Flowcytometry profiling demonstrated that CD4⁺ T cells from pancreatic tissue expressed higher frequencies of Glut-1⁺ and CD98⁺ cells than those from spleen or blood (Fig. 5c; ESM Fig. 5a, b), consistent with increased glucose and amino acid uptake in tissue-resident effector cells indicating cell activation, proliferation, and anabolic metabolism, while decreased expression suggests a resting or differentiated state. CD98 expression was high across compartments at diabetes onset, whereas CD71⁺ CD4⁺ T cells were more frequent and showed higher MFI in the spleen than in blood or pancreas (Fig. 5d; ESM Fig. 5a, b), suggesting ongoing proliferative expansion in secondary lymphoid organs. These data indicate that antigen encounter in this model is associated with both immune infiltration of the pancreas and tissue-specific metabolic activation of CD4⁺ T cells. Targeting these metabolic checkpoints may offer a strategy to selectively dampen pathogenic T-cell responses in early T1D [38–40].

**Fig. 5.**
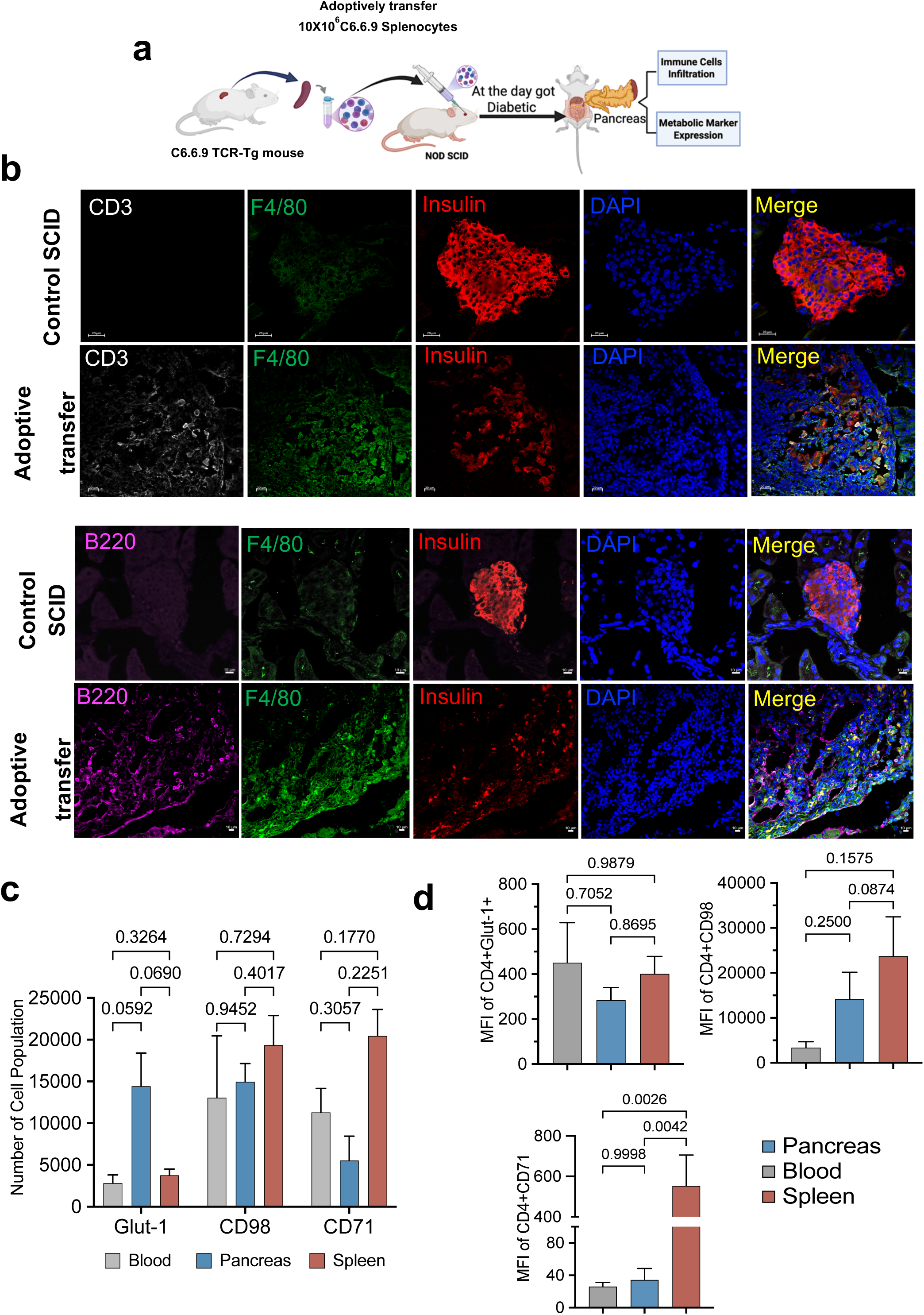
Metabolic profiling and immune infiltration in the pancreas following adoptive transfer of C6.6.9 T cells into NOD.Scid recipients. (**a**) Schematic overview of the experimental design used to assess metabolic status and immune infiltration in the pancreas after adoptive transfer. (**b**) Immunofluorescence staining of pancreatic sections from control and diabetic NOD.Scid mice, showing CD3^+^, F4/80^+^, and B220^+^ immune cell infiltration following BDC6.9 CD4⁺ T cell transfer. (**c, d**) Absolute number and MFI of metabolic markers (CD71, CD98, and Glut-1) expressed on CD4⁺ T cells isolated from adoptively transfer NOD.Scid mice peripheral blood, pancreas, and spleen at the time of diabetes onset. All data are presented as mean ± SEM (n = 4-10). Statistical significance was determined using one-way ANOVA with multiple comparisons, followed by Bonferroni post hoc test. P < 0.05 represents significant.

### Broadening and epitope diversification of IAPP-specific CD4⁺ T cells Co-recognizing ChgA and InsB9-23 antigen

To explore how antigen recognition evolves during disease, we re-analyzed adoptive transfer flow-cytometry data by gating on CD4⁺IAPP⁺ T cells and then assessing additional binding to ChgA and InsB9-23 tetramers over time (Fig. 6a; ESM Fig. 6a, b), This approach highlights the underutilized potential of BCR and TCR repertoire dynamics in diagnostic medicine. At baseline, very few CD4⁺IAPP⁺ cells co-stained with ChgA or InsB9-23 tetramers (Fig. 6b). After transfer into NOD.SCID recipients, the proportion and absolute number of CD4⁺IAPP⁺ChgA⁺ and CD4⁺IAPP⁺InsB9-23⁺ T cells in blood increased progressively and reached maximal levels around day 12 post-adoptive transfer (Fig. 6c, d). At diabetes onset, both dual positive subsets were enriched in pancreatic tissue compared with spleen (Fig. 6e, f). These findings indicate that within the IAPP-defined autoreactive pool, there is a broadening of tetramer-defined antigen recognition over time, with preferential accumulation of multi-positive cells in the pancreas during overt disease.

**Fig. 6.**
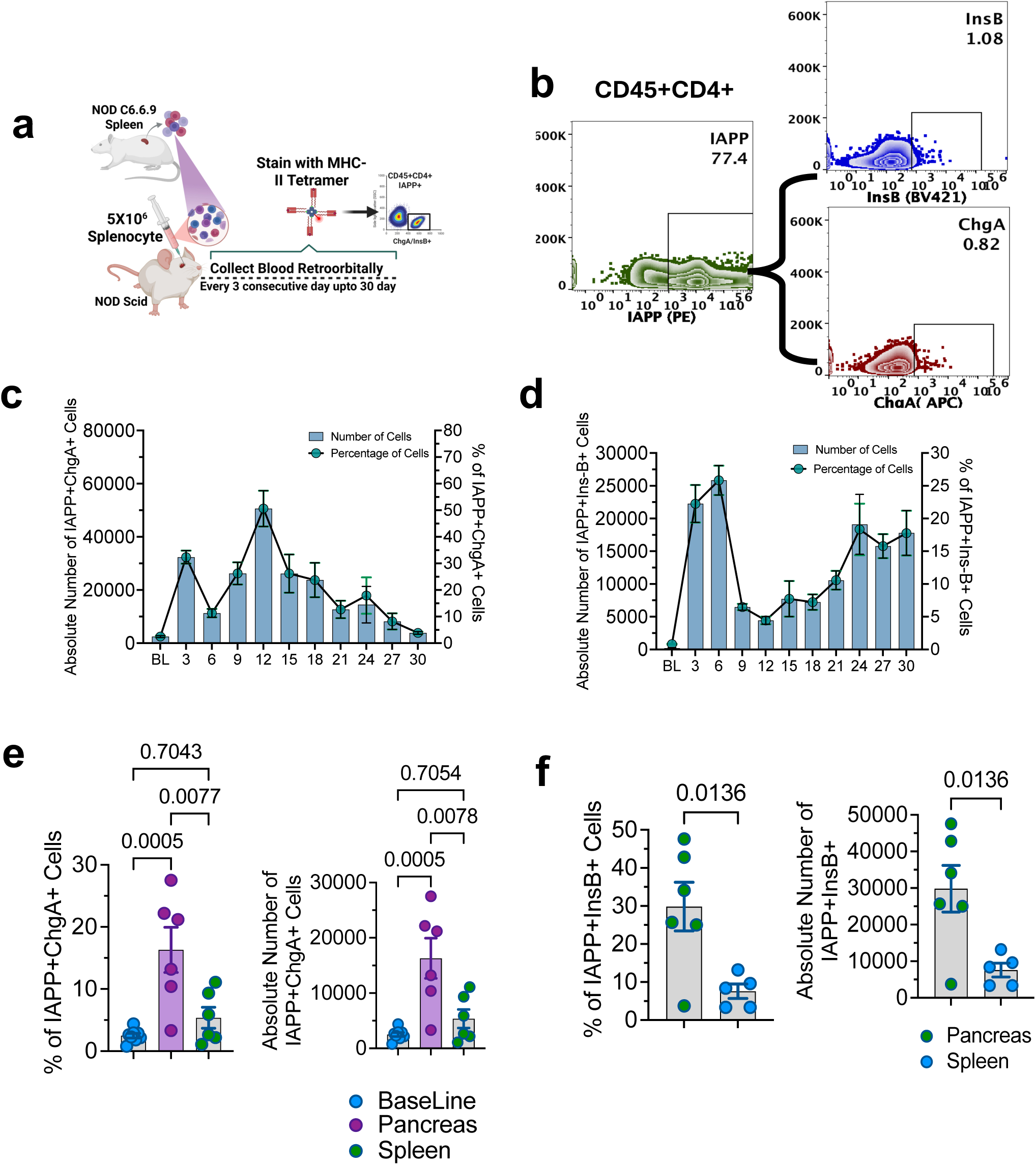
Phenotypic characterization of TCR rearrangement in IAPP-specific CD4⁺ T cells during diabetes progression in NOD.Scid recipient mice. (a) Schematic overview of experimental design for monitoring the epitope spreading following adoptive transfer of C6.6.9 TCR-Tg mouse CD4⁺ T cells into NOD.Scid recipient mice. (b) Densitometry flow cytometry plot representing baseline expression status of ChgA and InsB23-9 positive on C6.6.9 TCR-Tg mouse CD4⁺IAPP⁺ cells. (c) Frequency and absolute numbers of CD4⁺IAPP⁺ChgA⁺ and (d) CD4⁺IAPP⁺InsB9-23⁺ T cells in peripheral blood over time. (e, f) Frequency and absolute number of CD4⁺IAPP⁺ChgA⁺ and CD4⁺IAPP⁺InsB⁺ cells at the time of diabetes onset were assessed in pancreas, and spleen. All values are presented as mean ± SEM (n =5–6). Statistical significance was determined P < 0.05 represents significant using one-way ANOVA with multiple comparisons, followed by the Bonferroni post hoc test.

### sLAG-3 patterns in human T1D, FDRs, and longitudinal DEWIT cohorts

We then examined whether the sLAG-3 patterns observed in mice were recapitulated in human cohorts. In a cross-sectional analysis of individuals with type 1 diabetes up to six years after diagnosis and age-matched healthy controls, serum sLAG-3 levels did not differ significantly between groups (Fig. 7b). This suggests, as seen in the mouse experiments, that by the time of established disease, the early activation signal captured by sLAG-3 is not readily detectable. In contrast, when first-degree relatives were stratified by autoantibody status, autoantibody negative and single autoantibody FDRs displayed significantly higher sLAG-3 levels than multi-autoantibody FDRs, T1D subjects, and non-FDR-controls, indicating early T-cell activation (Fig. 7c). Consistent with this, sLAG-3 levels showed no significant differences among T1D patients regardless of autoantibody status (ESM Figure 7a), reinforcing that sLAG-3 is not a marker of late-stage disease burden.

**Fig. 7.**
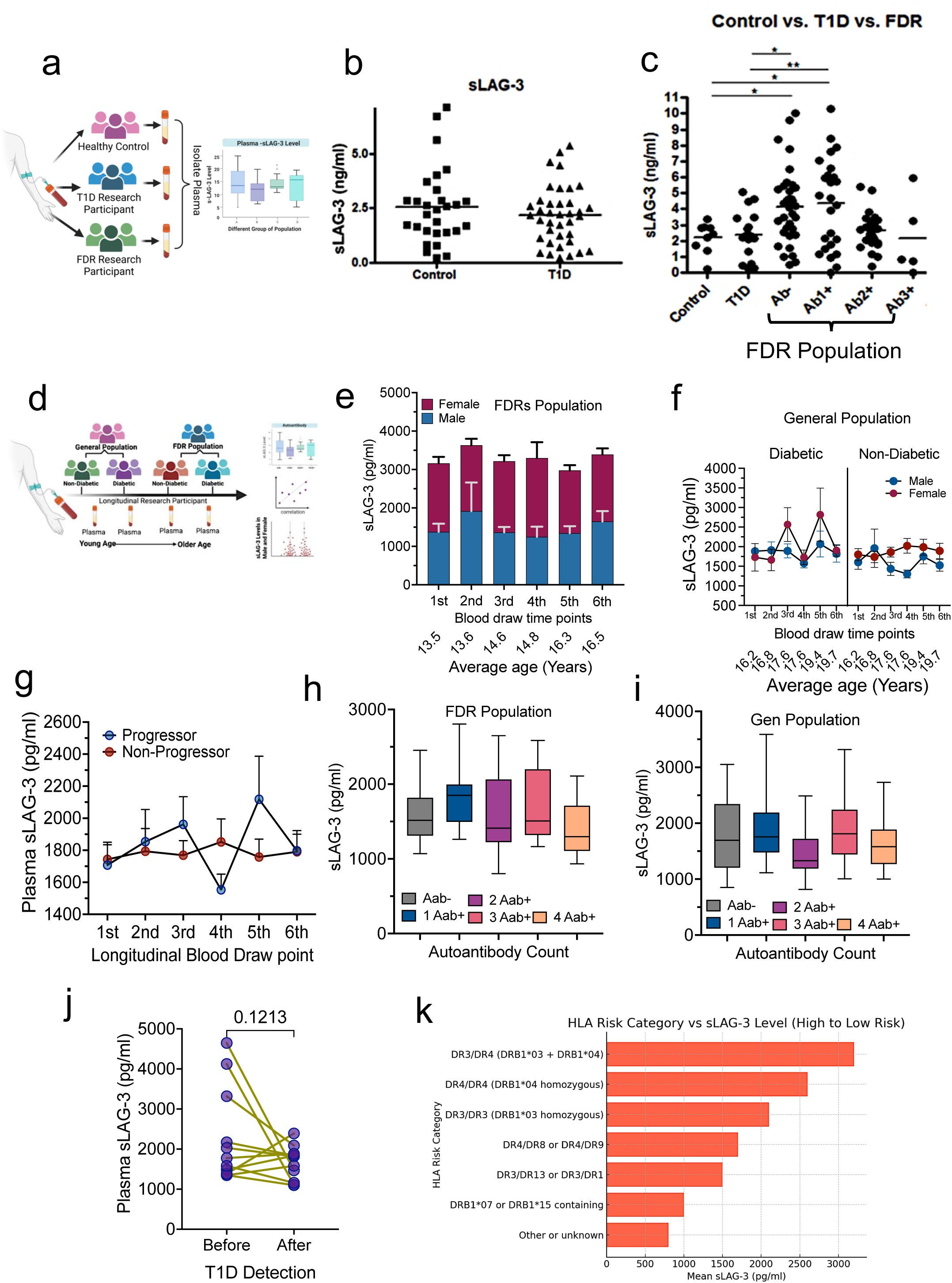
Longitudinal and cross-sectional analyses reveal early sLAG-3 elevation prior to autoantibody expansion and clinical onset of type 1 diabetes. (a) Experimental design for sample collection. Plasma was collected from healthy controls (n=28), type 1 diabetes patients (n=39), and FDRs with varying numbers of autoantibodies (Ab-, n=37 Ab1+, n=27; Ab2+, n=29; Ab3+, n=5). (b) Serum sLAG-3 levels in individuals with type 1 diabetes up to six years post-diagnosis was comparable to healthy controls, suggesting that heterologous immune responses may mask earlier elevations. (c) Autoantibody-negative and single-autoantibody FDRs show higher sLAG-3 levels than multiple-autoantibody individuals, type 1 diabetes subjects, and controls, indicating early T cell activation. (d) Experimental design for measurement of plasma sLAG-3 levels using MSD U-PLEX Assays in DEWIT and Stanford Cohorts. (e, f) DEWIT longitudinal cohort show higher sLAG-3 expression in females than males across both cohorts. (g) Progressor vs. non-progressor trajectories in the DEWIT cohort. Progressors display a gradual increase in sLAG-3 preceding diagnosis, whereas non-progressors show no change. (h, i) sLAG-3 levels stratified by autoantibody category in FDRs and the general population. (j) Longitudinal sLAG-3 profiling in DEWIT progressors shows a distinct peak immediately before diabetes onset followed by a sharp decline after diagnosis. (k) HLA genotype analysis demonstrating higher sLAG-3 expression in individuals carrying high-risk alleles. Statistical significance was determined using one-way ANOVA with multiple comparisons, followed by the Bonferroni post hoc test, *P < 0.05 significant.

Longitudinal analysis of the DEWIT cohort using MSD-based high sensitive assays, chosen for their suitability in repeated-measure analyses, showed that sLAG-3 levels were generally higher in females than males across both FDR and general population participants. (Fig. 7e, f). When participants were stratified by outcome, progressors exhibited a gradual rise in sLAG3 over time, culminating in a distinct peak just before diabetes diagnosis and a subsequent decline after onset. In contrast non-progressors showed no consistent change (Fig. 7g, j; ESM Fig. 7b). The sLAG-3 levels also tend-to decrease with increasing disease duration among individuals with type 1 dibetes (ESM Fig. 7c). In FDRs, sLAG-3 concentrations were highest in autoantibody negative and single-autoantibody positive individuals and progressively lower in those with multiple autoantibodies, with the single 4-autoantibody-positive individual showing the lowest value (Fig. 7h). This inverse relationship between sLAG-3 and autoantibody burden was not observed in general-population subjects (Fig. 7i), suggesting that early sLAG-3 elevation is most apparent in genetic or familial at-risk individuals. Additionally, non-diabetic individuals showed higher sLAG-3 concentrations compared with diabetic subjects among GADA positive, IA2A positive and ZnT8A positive participants in first-degree relatives, but no significant difference observed in general population. In contrast, IAA-positive groups showed no significant differences in sLAG-3 between diabetic and non-diabetic individuals, indicating that alterations in circulating sLAG-3 are associated with diabetes status in specific autoantibody-defined subgroups (ESM Fig. 7d-g). Finally, sLAG-3 levels were higher in subjects carrying high risk HLA alleles such as DRB1\**04* (DR4) *and DQA1\**03:01-DQB1*03:02 (DQ8) compared with those lacking these risk genotypes (Fig. 7k). This result support a linking between genetic susceptibility and the magnitude of early immune activation.

### sLAG-3 dynamics during COVID-19 vaccination and CVB infection

To test whether sLAG-3 reliably marks early T cell activation in nonautoimmune contexts, we measured sLAG-3 following COVID-19 vaccination in healthy adults and during Coxsackievirus B3 (CVB3) infection in mice. In vaccinated adults, plasma sLAG-3 (measured by Olink as NPX values) increased significantly by 10 days after vaccination compared with pre-vaccination levels, with most donors showing a clear upward shift (Fig. 8a, b). This transient increase occurred in the expected window for vaccine induced T cell responses and before the mature humoral response.

**Fig. 8.**
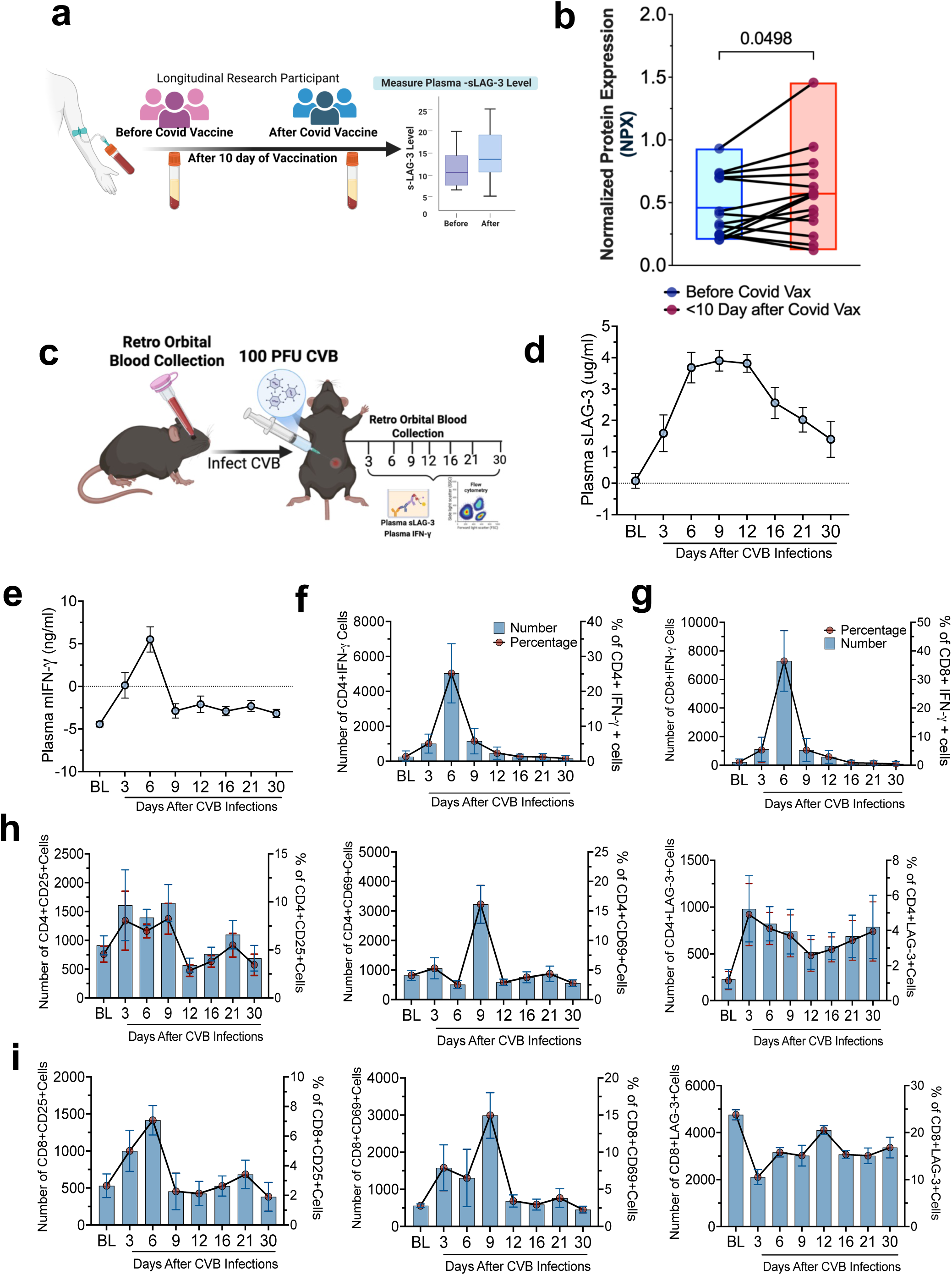
sLAG-3 as an early and sensitive marker of T cell activation during vaccination and viral infection C57BL/6 Mice. (a) Schematic illustration of before and 10 days after COVID-19 vaccination, indicating rapid T-cell activation preceding antibody development. (b) Normalized protein expression (NPX) of sLAG-3 over time in before and after 10 days of Covid vaccination from plasma. Each donors’ samples are connected by a line. (c) Schematic representation of time-course analysis of plasma analytes in-vivo CVB-infected C57BL/6 mice. (d) Plasma sLAG-3 levels form C57BL/6 mice infected with CVB. (e) Plasma IFN-γ concentrations increase after infection, with a marked rise at day 6, reflecting concurrent immune activation. (f, g) Flow cytometry reveals time-dependent increases in IFN-γ-producing CD4⁺ and CD8⁺ T cells in peripheral blood following CVB infection. (h, I) Expression of T cell activation markers CD25 and CD69. All values are presented as mean ± SEM (n=8). Statistical significance was determined using one-way ANOVA, followed by the Bonferroni post hoc test. *P < 0.05 represent significant.

In CVB infected C57BL/6 mice, plasma sLAG-3 was minimal at baseline, start to increase by day 3, and peaked between days 6-9 before declining toward baseline by day 30 (Fig. 8c, d). Plasma IFN-γ levels followed a similar pattern with a marked increase around day 6 (Fig. 8e). Flow cytometry revealed a concurrent expansion of IFNγ-producing CD4⁺ and CD8⁺ T cells and increased expression of activation markers CD25 and CD69 in peripheral blood (Fig. 8f-i; ESM Fig. 8). These data confirm that sLAG-3 is a sensitive and dynamic marker of early T-cell activation in both vaccine and viral infection settings.

## Discussion

Our goal was to determine whether soluble LAG-3 is useful serve as a biomarker of early T cell activation relevant to type 1 diabetes pathogenesis. Across spontaneous and adoptive transfer NOD mouse models, human FDR and longitudinal type 1 diabetes cohorts, and independent vaccine and viral infection systems. We consistently found that sLAG-3 concentrations in plasma/serum rises significantly early during T cell activation and declines as responses contract or as overt autoimmune diabetes develops. These observations support sLAG-3 as a candidate biomarker of early T cell activation and warrant further evaluation of its utility for staging autoimmune risk [16].

In NOD mice, plasma sLAG-3 increased between 6-12 weeks of age and returned to near baseline by 16 weeks, while blood glucose remained normal until around 16 weeks of age, when overt diabetes onset occurred. Consistent with our findings, Mathews and Ludlow, demonstrated that subtle blood glucose dysregulation precedes overt diabetes and extend extends this work by providing an immunological basis for these early changes [41]. Their results suggest that early blood glucose elevations reflect increased antigen-specific, immune-mediated beta-cell stress driven in part by BDC-2.5 reactive CD4⁺ T cells rather than global beta-cell loss, positioning early blood glucose perturbations as a functional marker of immune activation preceding clinical diabetes [41]. Interestingly, we found that, beta-cell antigen specific CD4⁺ T cells defined by InsB923, ChgA, and IAPP tetramers expanded over the same 6-12-week window in the same animal cohort. These findings correlated positively with sLAG-3 plasma concentrations. Furthermore, in the C6.6.9 adoptive transfer model, sLAG-3 and *Ins2* mRNA rose rapidly after adoptive transfer that peaked well before hyperglycemia. Histological and flow cytometric evidence of insulitis and donor CD4⁺ T cell accumulation in the pancreas. Together, these results indicate that sLAG-3 is induced and persists during early antigen-driven activation of autoreactive CD4⁺ T cells and that its peak precedes frank metabolic decompensation [42].

The adoptive transfer experiments also delineate how activation, tissue localization, and metabolism among autoreactive CD4⁺ T cells evolve during disease progression. CD25 and CD69 expression on peripheral CD4⁺ T cells peaked early after transfer, paralleling the sLAG-3 surge, and our detection of tetramer-positive cells. Subsequently, T cells accumulated in the pancreas, where CD69 expression increases. CD4⁺ T cells, that had infiltrated the pancreas exhibit increased expression of Glut-1, CD98, and CD71 expression relative to blood and spleen, consistent with metabolic reprogramming of effector cells in the inflamed target organ [43–49]. Within the IAPP specific pool, the emergence and pancreatic localization of CD4⁺IAPP⁺ChgA⁺ and CD4⁺ IAPP⁺ InsB2-93⁺ subsets during diabetes indicated a progressive broadening of tetramer-defined antigen recognition over time. Our results provide a coherent picture in which early antigen encounter drives a transient systemic sLAG-3 signal, expansion and tissue homing of antigen-specific CD4⁺ T-cells and increased metabolic activity in the pancreas, long before elevated blood glucose and failure of first phase insulin loss [33].

In humans, the cross sectional T1D vs non-diabetic control comparison showed no difference in sLAG-3 levels up to six years post diagnosis, implying that the early activation window identified in mice is not captured when sampling individuals with established disease. In contrast, FDRs at earlier disease stages displayed a distinct pattern: autoantibody negative and single autoantibody FDRs had higher sLAG-3 than multi-autoantibody positive FDRs, T1D subjects, and non-FDR controls. Thus sLAG-3 declined with increasing autoantibody burden. Longitudinal data from the DEWIT cohort further showed that progressors, but not non-progressors, exhibited a gradual rise in sLAG-3 culminating in a peak immediately prior to diagnosis that declined thereafter. sLAG-3 tended to be higher in individuals carrying higher risk type 1 diabetes HLA genotypes. Taken together, these human data are consistent with a model in which sLAG-3 in blood serving as a biomarker of autoimmunity during a relatively narrow early stage of immune activation that precedes or is co-incident with a broadening of autoantibody and auto-reactive T cell expansion prior to clinical onset in genetically susceptible individuals [50, 51].

The current human dataset has important limitations that temper how strongly sLAG-3 can be positioned as a “surrogate marker before antibodies.” First, most of the human analyses are cross-sectional. Even in the DEWIT cohort, the number of progressors with dense longitudinal sampling is limited in. numbers. We therefore cannot yet systematically align the timing of sLAG-3 concentration with individual seroconversion events to prove that sLAG-3 consistently rises before the first islet autoantibody appears. Second, sLAG-3 levels were measured using different assay platforms and in relatively small subgroups, which constrains our ability to define robust thresholds or evaluate test characteristics such as sensitivity, specificity, and predictive value. Third, the COVID-19 vaccine and CVB infection experiments highlight that increases in sLAG-3 in blood occurs general following T cell activation and may not be specific to autoimmunity. Eelevations in at-risk individuals will likely need to be interpreted in the context of recent infections, vaccinations, or other immune stimuli.

The COVID19 vaccination and CVB infection data nonetheless provide important context. In both settings, sLAG-3 rose rapidly during the effector phase of the immune response, coincided with increases in IFN-γ and T-cell activation but markers and returned to baseline as the response resolved. These observations support the idea that sLAG-3 is a sensitive and reversible readout of early T-cell activation. In type 1 diabetes, this suggests that transient sLAG-3 elevations in FDRs are biologically plausible and that serial measurements may be more informative than single time points, especially when integrated with autoantibody profiles, islet or beta-cell antigen–specific T cell along with detection of biorepositories, and plasma circulating insulin mRNA assessments in the clinical context provide additional important data that may help distinguish autoimmunity from other T cell-mediated immune responses.

Overall, our findings support as a cautious interpretation that sLAG-3 is a promising biomarker of early T cell activation, detectable in some autoantibody-negative and single autoantibody positive first-degree relatives and declining with increasing autoantibody burden and after diabetes onset. When integrated with beta-cell antigen specific tetramer profiling and markers of beta-cell stress such as circulating *Ins2* mRNA, sLAG-3 may help define an early immunological phase characterized by autoreactive CD4⁺ T cell activation can be detected before substantial beta-cell loss is clinically evident. Larger, deeply phenotype longitudinal cohorts will need to be tested to determine if assessment of sLAG-3, autoantibody reactivity, markers of beta cell stress (circulating insulin mRNA in blood) and T cell phenotypes comprise surrogate markers of type 1 diabetes risk or progression and whether these markers are predictive of patients that go on to develop type 1 diabetes clinically to determine how frequently it precedes seroconversion at the individual level.

## Conclusion

In conclusion, our findings support sLAG-3 as a candidate early biomarker of T cell activation in type 1 diabetes, with evidence that sLAG-3 may be elevated before or at the earliest stages of islet autoimmunity in individual development in some at-risk individuals. We propose that tracking of beta-cell antigen specific CD4⁺ T cell, circulating *Ins2* mRNA measurements, metabolic profiling, and analyses of B and T cell epitope spreading in conjunction with sLAG-3 reflects an early immunological state associated with autoreactive T-cell diversification rather than late-stage beta-cell loss. Nevertheless, larger longitudinal studies aligned to seroconversion are required to validate sLAG-3 as a surrogate marker of pre-autoantibody autoimmune disease and to define its utility for early risk stratification and therapeutic intervention.

## Data and Resource Availability

The data sets generated and/or analyzed during the current study are available from the corresponding author on reasonable request.

## Acknowledgements

The authors thank Dr. Dario Vignali lab at University of Pittsburgh for providing the secondary antibody for the mouse sLAG-3 ELISA. We also thank to Dale Long - Facility Manager at NIH Tetramer Core Facility at Emory for providing tetramers and the Center for Diabetes and Metabolic Diseases for access to its instrumental facilities.

## Funding

This study was funded by the National Institute of Diabetes and Digestive and Kidney Diseases (NIDDK) under project 7R01DK132583, and Breakthrough Type 1 diabetes research Strategic Research Agreement (SRA) funds, under Project 1-PNF-2025-1642-A-N to Dr. Jon D. Piganelli, and R01DK129287 to Jason M. Spaeth. Farooq Syed was supported by a Breakthrough T1D (formerly JDRF) CDA (5-CDA-2022-1176-A-N and (CDA)/3-CDA-2025-1692-A-N) and Thomas Beatson Foundation (2024-010). This study is also supported by a NIH-R21 (1R21AI193399) award to Jon D. Piganelli and Farooq Syed.

## Contribution statement

J.D.P., S.R., F.S., and C.S. conceptualized the study. S.R. and J.D.P. designed all experiments; S.R. and M.L.P. performed experiments. S.R. analyzes all data and prepares all figures. F.S and J.R performed experiments on insulin mRNA, and M.E.S. conducted sLAG-3 assays in Covid-vaccinated participants. S. R and A.D.S. analyzed all confocal images. W.H help to collect the DEWIT cohort samples, A.T., E.S.H., E.M., and K.P.J. collect human plasma sample from Stanford Biobank, J.S. contributed to discussions and study design. S.R. and J.D.P. drafted the manuscript. S.F., M.L.P., J.S., A.D.S., K.P.J., J.S., and C.S. editing and finalizing it. All authors have no conflicts of interest.

## Abbreviations

ChgA: Chromogranin A
FACS: Fluorescence-Activated Cell Sorting buffer
FDR: first degree relative
GAD65: Glutamic Acid Decarboxylase 65 (isoform of GAD)
HLA: Human Leukocyte Antigen
IA-2: Islet Antigen-2 (a pancreatic β-cell antigen)
IAA: Insulin Autoantibody
IAPP: Islet Amyloid Polypeptide (also called amylin)
IFN-γ: Interferon-gamma
mRNA: messenger Ribonucleic Acid
NOD: Non-Obese Diabetic (mouse strain)
NOD.SCID: NOD. Severe combined immunodeficient
RPM: Revolutions Per Minute (common lab term, e.g., centrifugation speed)
TCR: T Cell Receptor
ZnT8: Zinc Transporter 8

## Notes

### Competing Interest Statement

The authors have declared no competing interest.

### Summary of Updates

We would like to inform you that there was a mistake in the spelling of the author name, Jason Spaeth. The correction has now been made. We kindly request that the necessary updates be applied accordingly to reflect this change. Thank you for your assistance

