## Supplementary Figure for "Soluble LAG-3 Identifies a Dynamic Early T Cell Activation Window in self-reactivity, Type 1 Diabetes, and Broader Immune Responses"

***Corresponding Authors:**

[a] Both authors equally contribute.

ESM Figure 1.

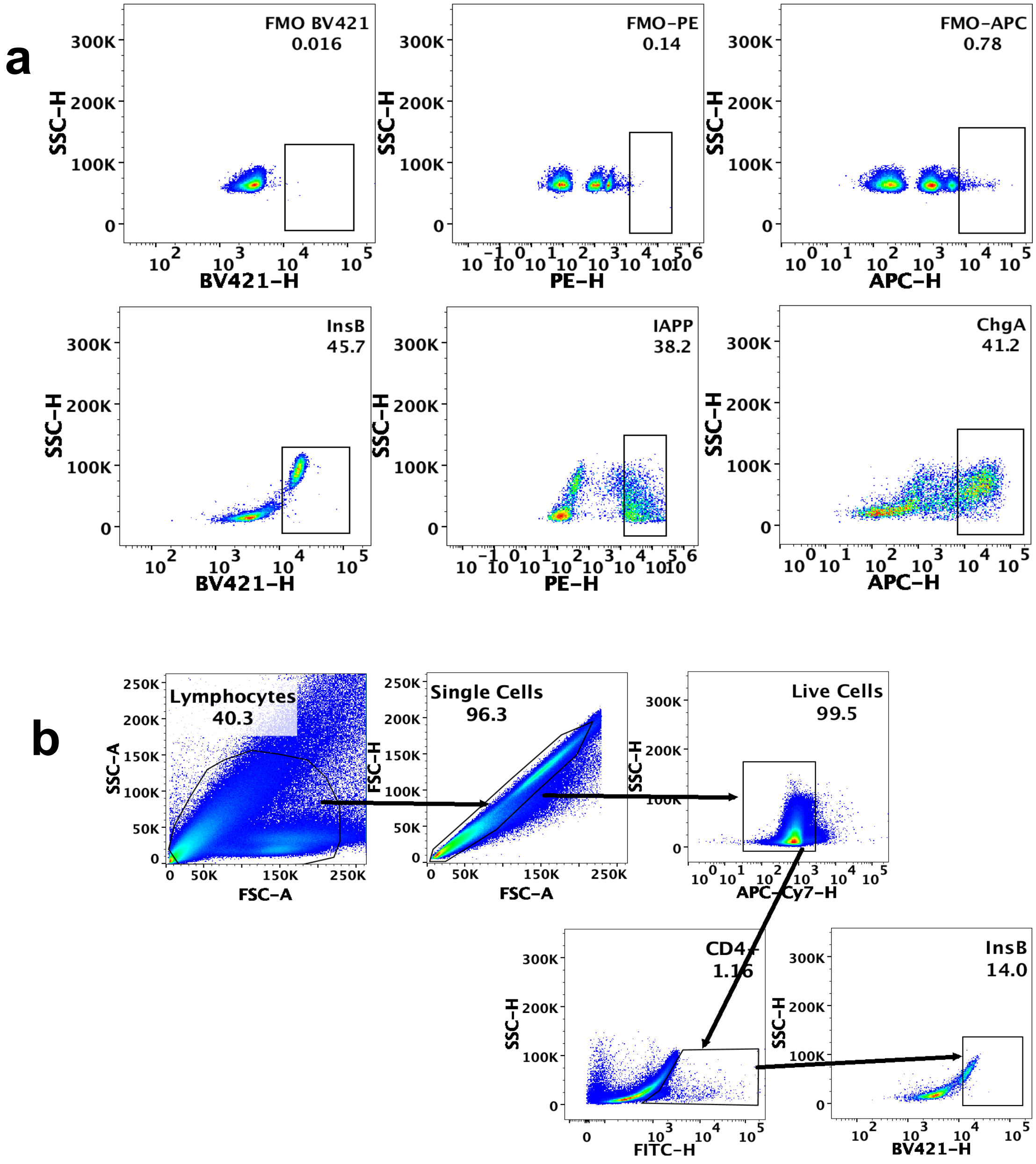


**ESM Figure 1.** Flow cytometry gating strategies. (**a**) Fluorescence minus one (FMO) Gating strategy used to define activation marker expression on tetramer-positive $CD4^+$ T cells. (**b**) Sequential gating of lymphocytes by forward scatter (FSC), side scatter (SSC), followed by live-cell, and $CD4^+$ T cell gates to identify tetramer-positive populations.

ESM Figure. 2

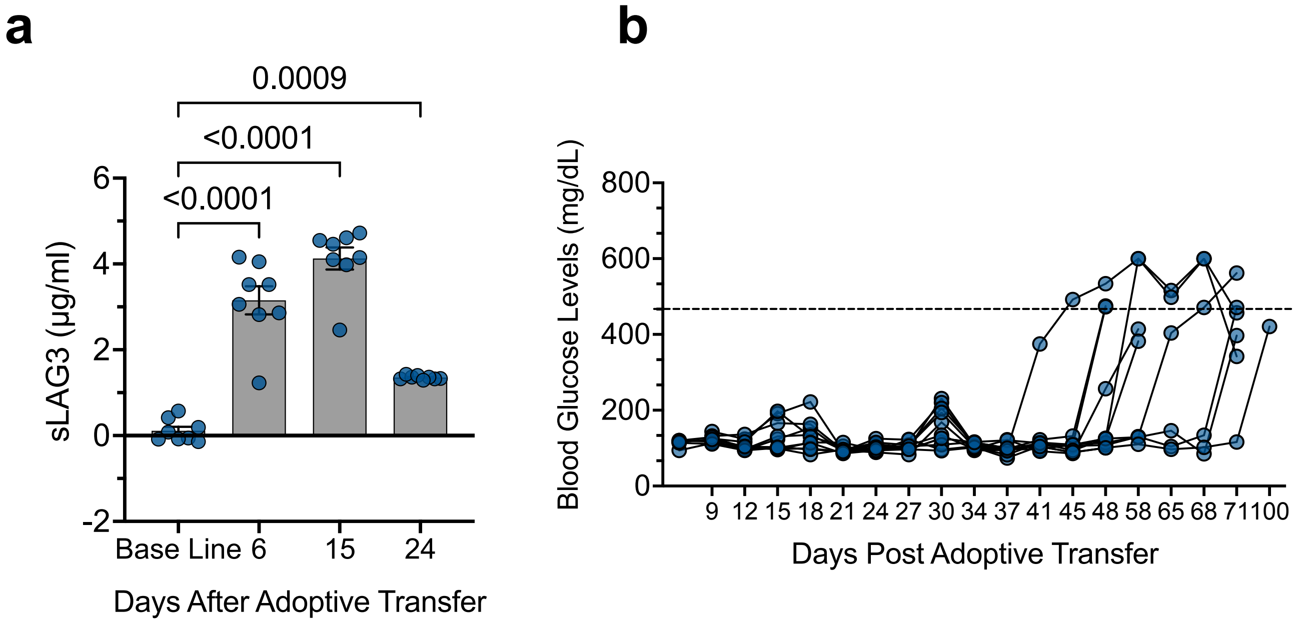


**ESM Figure 2.** Plasma sLAG-3 dynamics following post-transfer. (**a**) Quantification of plasma sLAG-3 levels at indicated time points following adoptive transfer. (**b**) Individual blood glucose trajectories of adoptively transferred NOD.SCID mice. Data are presented as mean ± SEM (n=10). Statistical significance was determined using one-way ANOVA with Bonferroni post-hoc testing; *$p<0.05$.

ESM Figure 3.

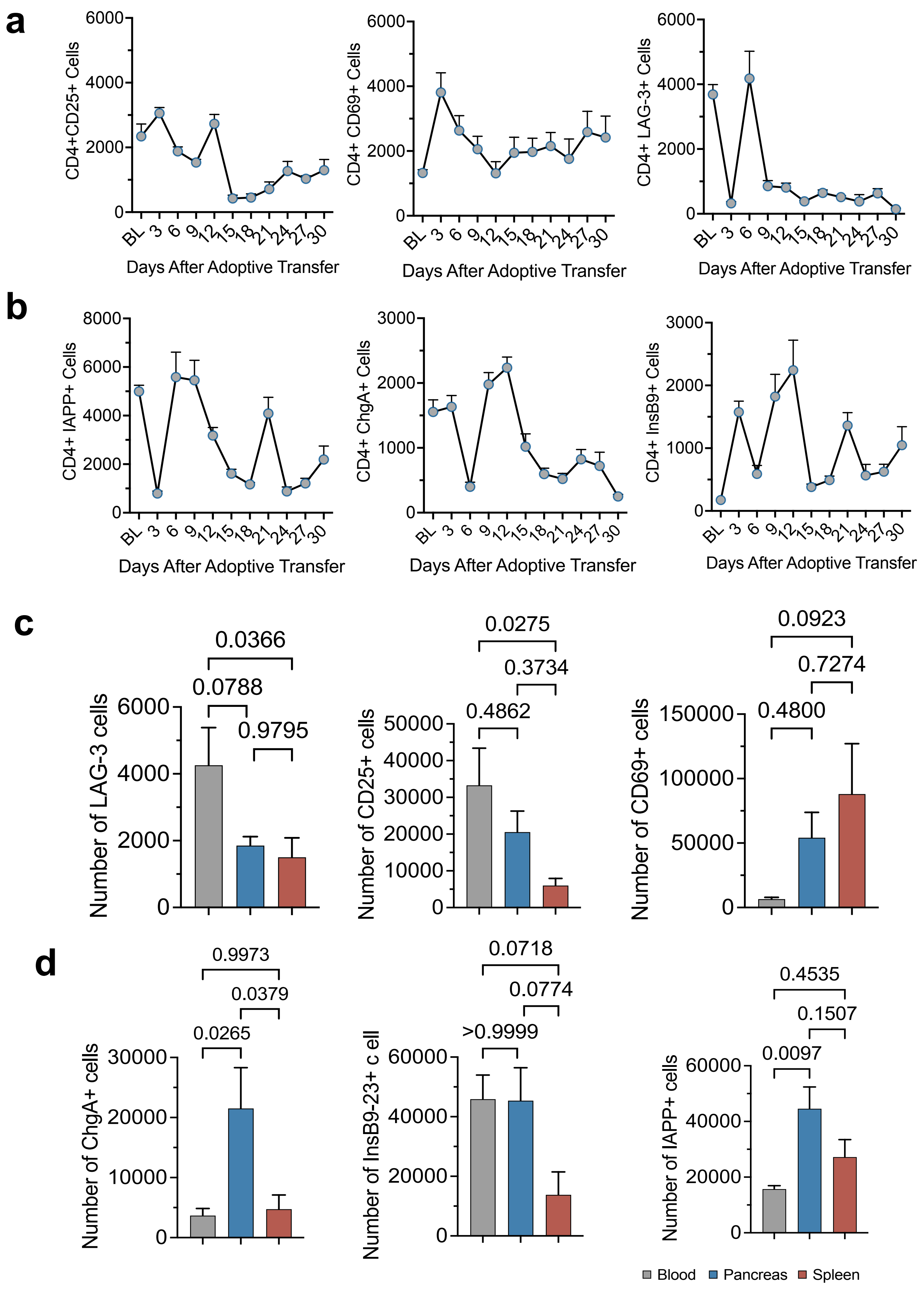
a
CD4+CD25+ Cells
CD4+ CD69+ Cells
CD4+ LAG-3+ Cells
Days After Adoptive Transfer
BL 3 6 9 12 15 18 21 24 27 30
b
CD4+ IAPP+ Cells
CD4+ ChgA+ Cells
CD4+ InsB9+ Cells
Days After Adoptive Transfer
c
Number of LAG-3 cells
0.0366
0.0788
0.9795
Number of CD25+ cells
0.0275
0.3734
0.4862
Number of CD69+ cells
0.0923
0.7274
0.4800
d
Number of ChgA+ cells
0.9973
0.0379
0.0265
Number of InsB9-23+ c ell
0.0718
0.0774
>0.9999
Number of IAPP+ cells
0.4535
0.1507
0.0097
Blood
Pancreas
Spleen

**ESM Figure 3.** Phenotypic characterization of T cell markers in adoptively transferred C6.6.9 T-Cells. (**a**) Absolute numbers of $CD25^+$, $CD69^+$, and LAG-$3^+$ $CD4^+$ T cells in peripheral blood over time following adoptive transfer. (**b**) Absolute numbers of β-cell antigen-specific tetramer-positive $CD4^+$ T cells (ChgA, InsB9-23, IAPP) in peripheral blood post-transfer. (**c, d**) Distribution of activation markers and tetramer-positive $CD4^+$ T cells in blood, pancreas, and spleen at time of diabetes onset. Data are presented as mean ± SEM (n=4-10). Statistical significance was determined using one-way ANOVA with Bonferroni post-hoc testing; *$p<0.05$.

ESM Figure 4.

**a** Baseline profiling of blood lymphocytes in NOD-SCID mice before adoptive transfer.

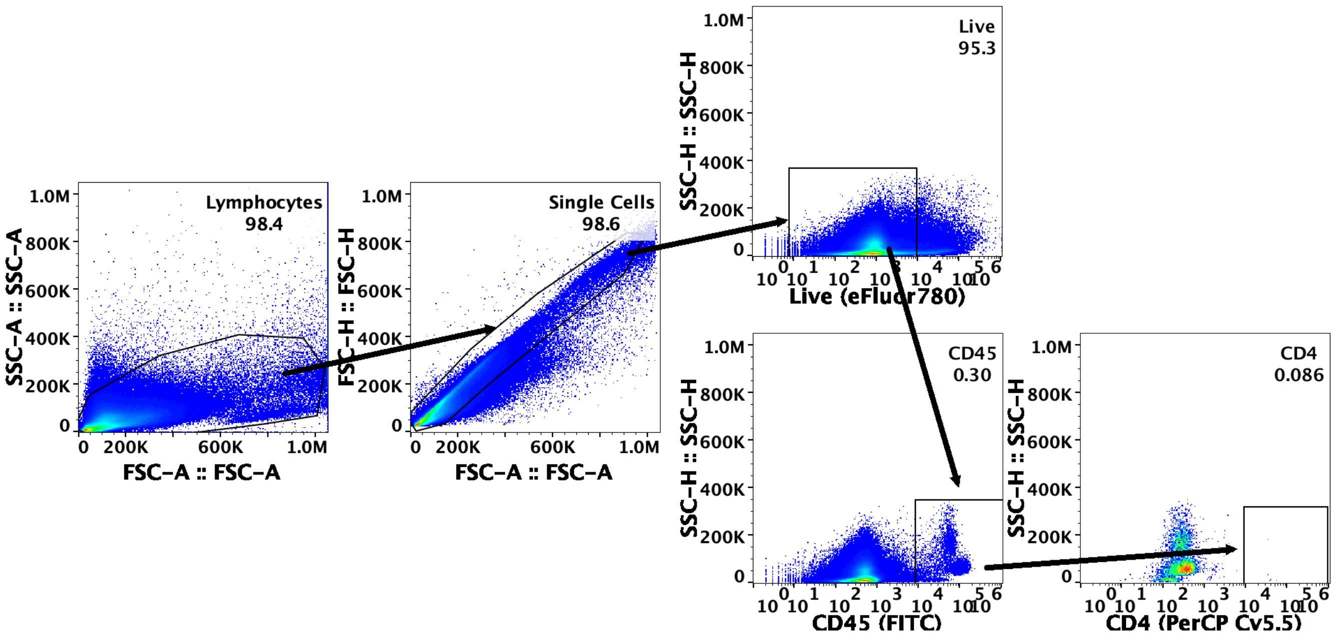


**b** Gating strategy for surface markers and Tetramer expression in BDC6.9 Spleenocyte

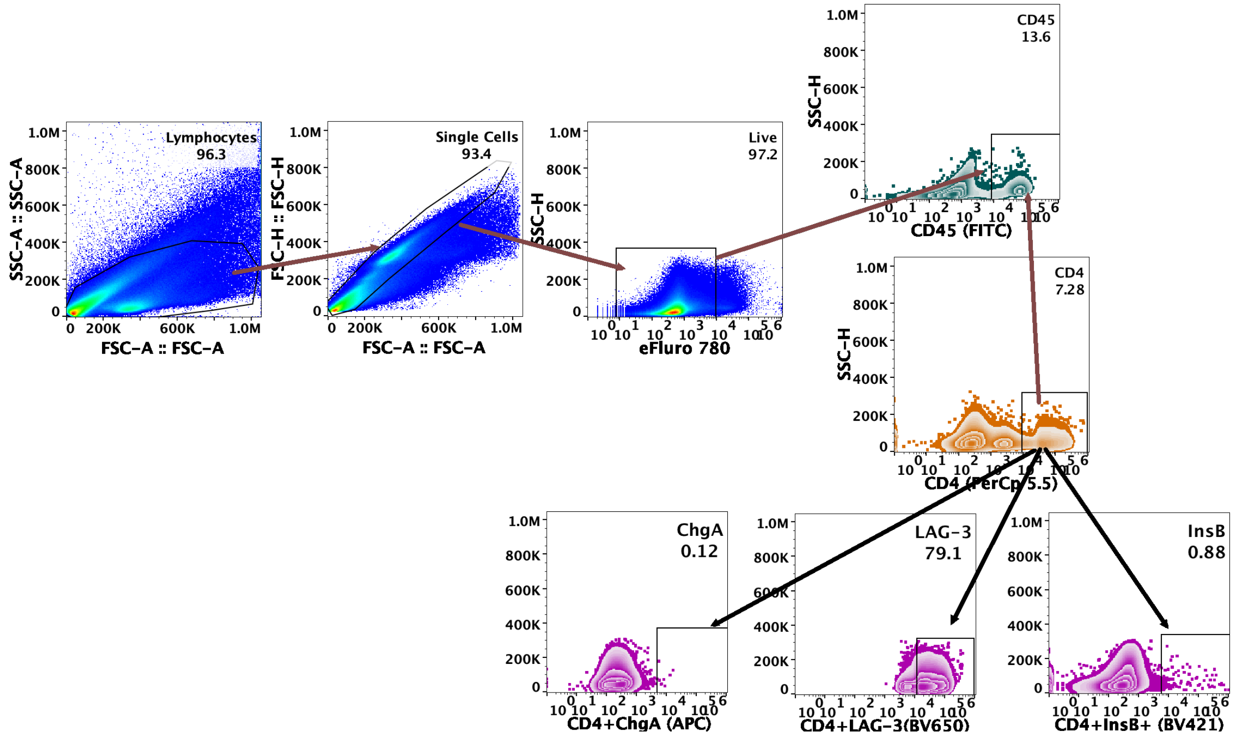

**ESM Figure 4.** Baseline and post-transfer gating strategies. (**a**) Baseline expression of different surface markers on peripheral blood lymphocytes in NOD.SCID mice prior to adoptive transfer. (**b**) Flow cytometry gating strategy used to assess surface marker and tetramer expression on $CD4^{+}$ T cells following adoptive transfer.

ESM Figure 5.

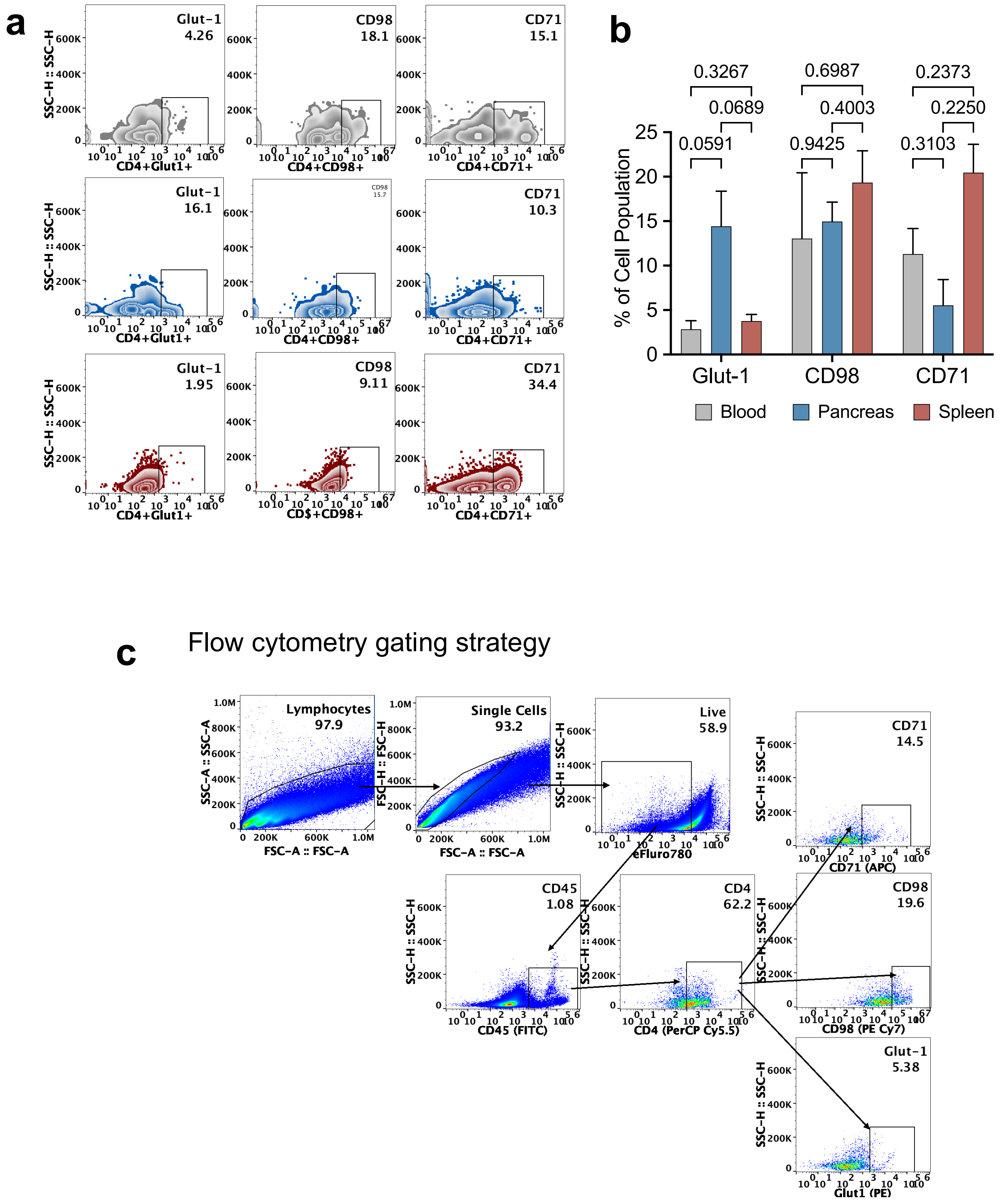

a
Glut-1
4.26
CD98
18.1
CD71
15.1
SSC-H :: SSC-H
CD4+Glut1+
CD4+CD98+
CD4+CD71+
Glut-1
16.1
CD98
15.7
CD71
10.3
Glut-1
1.95
CD98
9.11
CD71
34.4
CD$+CD98+
b
0.3267
0.6987
0.2373
0.0689
0.4003
0.2250
0.0591
0.9425
0.3103
% of Cell Population
Glut-1
CD98
CD71
Blood
Pancreas
Spleen
c
Flow cytometry gating strategy
Lymphocytes
97.9
Single Cells
93.2
Live
58.9
CD71
14.5
SSC-A :: SSC-A
FSC-A :: FSC-A
FSC-H :: FSC-H
eFluro780
CD71 (APC)
CD45
1.08
CD4
62.2
CD98
19.6
CD45 (FITC)
CD4 (PerCP Cy5.5)
CD98 (PE Cy7)
Glut-1
5.38
Glut1 (PE)

**ESM Figure 5.** Expression of T-cell metabolic markers at diabetes onset. (**a**) Representative density plots and (**b**) Quantitative bar graphs showing the percentage of $CD4^+$ T-cells expressing metabolic markers (CD71, CD98, and Glut-1) in peripheral blood, pancreas, and spleen at the time of diabetes onset. (**c**) Flow cytometry gating strategy for metabolic marker analysis. Lymphocytes were first gated on forward scatter (FSC) and side scatter (SSC), followed by gating on live $CD45^+CD4^+$ T cells to assess the expression of CD71, CD98, and Glut-1. Data are presented as mean ± SEM (n=5). Statistical significance was determined using one-way ANOVA with Bonferroni post-hoc testing; $*p<0.05$.

ESM Figure 6.

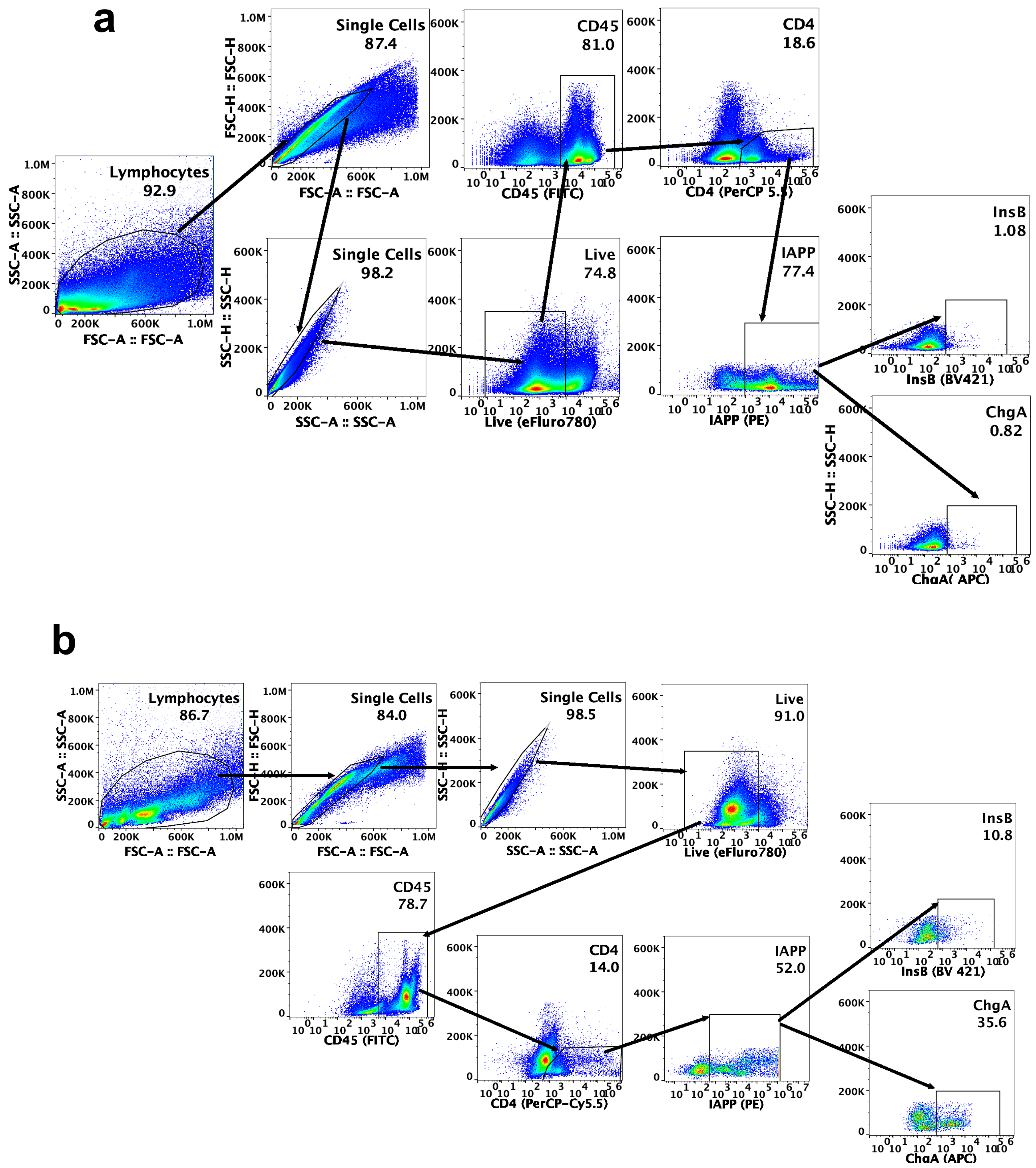
a
Lymphocytes
92.9
SSC-A :: SSC-A
FSC-A :: FSC-A
Single Cells
87.4
FSC-H :: FSC-H
FSC-A :: FSC-A
Single Cells
98.2
SSC-H :: SSC-H
SSC-A :: SSC-A
CD45
81.0
CD45 (FITC)
Live
74.8
Live (eFluro780)
CD4
18.6
CD4 (PerCP 5.5)
IAPP
77.4
IAPP (PE)
InsB
1.08
InsB (BV421)
ChgA
0.82
SSC-H :: SSC-H
ChgA( APC)
b
Lymphocytes
86.7
SSC-A :: SSC-A
FSC-A :: FSC-A
Single Cells
84.0
FSC-H :: FSC-H
FSC-A :: FSC-A
Single Cells
98.5
SSC-H :: SSC-H
SSC-A :: SSC-A
Live
91.0
Live (eFluro780)
CD45
78.7
CD45 (FITC)
CD4
14.0
CD4 (PerCP-Cy5.5)
IAPP
52.0
IAPP (PE)
InsB
10.8
InsB (BV 421)
ChgA
35.6
ChgA (APC)

**ESM Figure 6.** Flow cytometry gating strategy used to analyze Phenotypic characterization of IAPP$^+$ T-cell receptor rearrangement during diabetes progression in adoptively transferred NOD.SCID mice Flow cytometry gating strategy for epitope spreading analysis. (**a**) Baseline gating of C6.6.9 splenocytes before adoptive transfer of C6.6.9 T cells into NOD.SCID mice. (**b**) Post-transfer gating to identify CD45$^+$CD4$^+$IAPP$^+$ T cells, while assessing their ChgA and InsB9-23 reactivity over diabetes progression. Lymphocytes were first gated on FSC and SSC, followed by gating on live CD45$^+$CD4$^+$IAPP$^+$ T cells to assess the expression of InsB9$^+$ and ChgA$^+$ cells population.

ESM Figure 7.

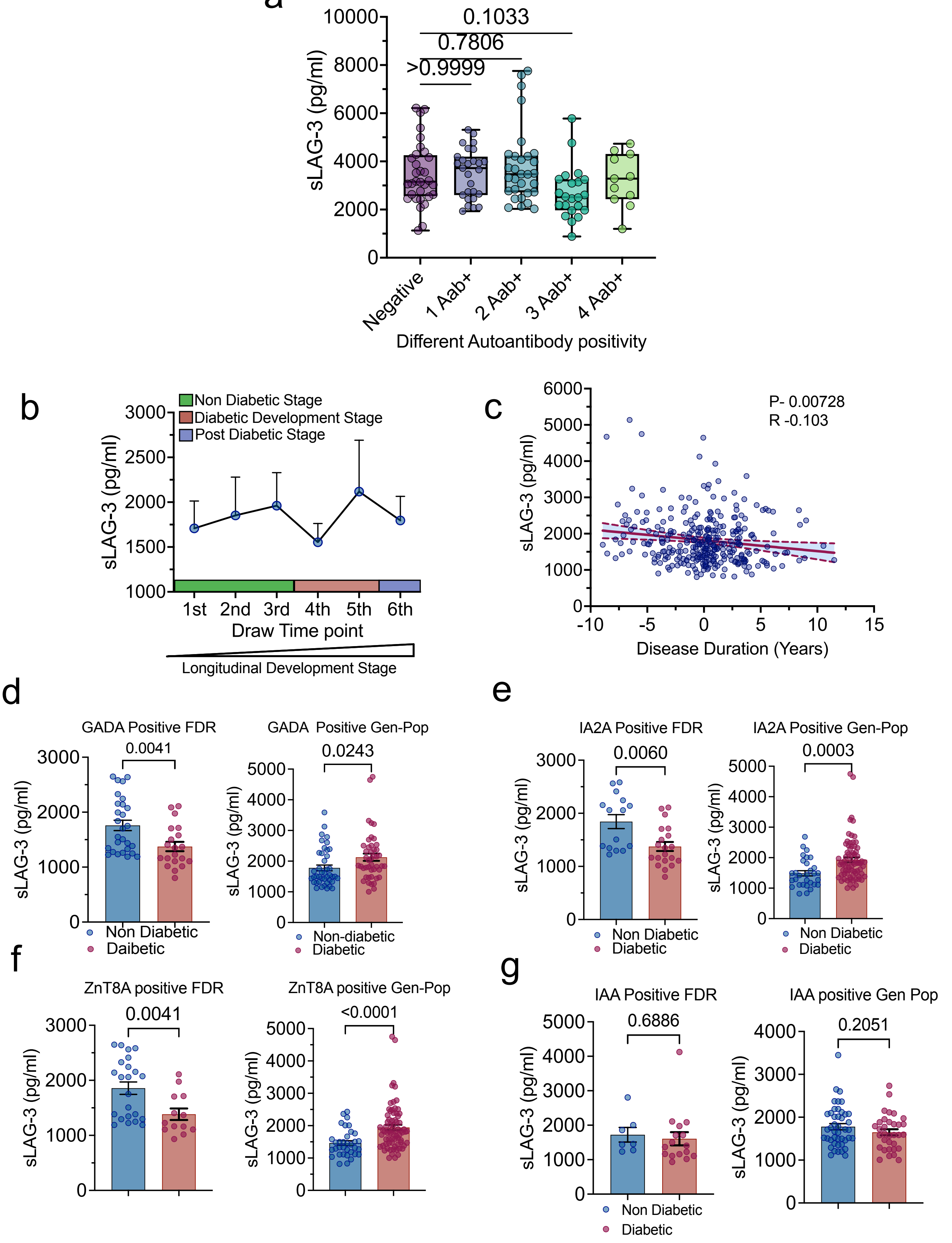
a
0.1033
0.7806
>0.9999
sLAG-3 (pg/ml)
Negative
1 Aab+
2 Aab+
3 Aab+
4 Aab+
Different Autoantibody positivity
b
Non Diabetic Stage
Diabetic Development Stage
Post Diabetic Stage
sLAG-3 (pg/ml)
1st
2nd
3rd
4th
5th
6th
Draw Time point
Longitudinal Development Stage
c
P- 0.00728
R -0.103
sLAG-3 (pg/ml)
Disease Duration (Years)
d
GADA Positive FDR
0.0041
GADA Positive Gen-Pop
0.0243
Non Diabetic
Daibetic
Non-diabetic
Diabetic
e
IA2A Positive FDR
0.0060
IA2A Positive Gen-Pop
0.0003
Non Diabetic
Diabetic
f
ZnT8A positive FDR
0.0041
ZnT8A positive Gen-Pop
<0.0001
g
IAA Positive FDR
0.6886
IAA positive Gen Pop
0.2051
Non Diabetic
Diabetic

**ESM Figure 7**. Cross-sectional and longitudinal analysis of plasma sLAG-3 levels by autoantibody status, disease duration, and diabetes stage. (**a**). Plasma sLAG-3 levels in diabetic individuals, grouped by autoantibody status (Aab-, 1Aab+, 2Aab+, 3Aab+, 4Aab+). (**b**) Longitudinal sLAG-3 profiles from DEWIT samples, across different developmental stages (non-diabetic, diabetic-development, and post-diabetic) showing variable but stage-dependent fluctuations. (**c**) Negative correlation between plasma sLAG-3 levels and disease duration in T1D subjects. (**d-g**) sLAG-3 concentrations in GADA, IA2A, ZnT8A and IAA positive individuals from FDR and general-population cohorts, comparing diabetic and non-diabetic subjects. All values are presented as mean ± SEM. Statistical significance was determined $P < 0.05$ represents significant using t-test.

### ESM Figure 8.

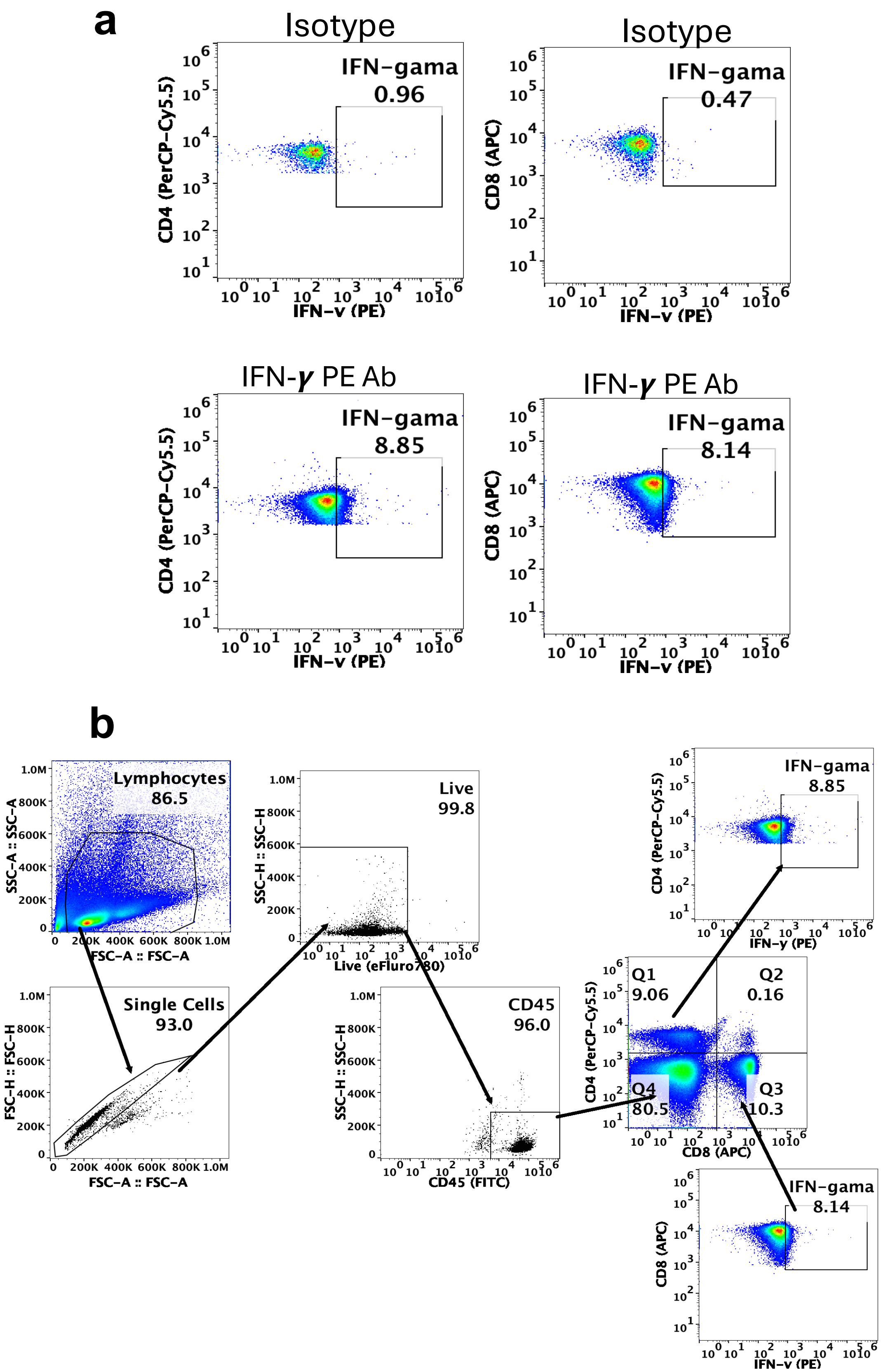

a
Isotype
IFN-gama
0.96
CD4 (PerCP-Cy5.5)
IFN-γ (PE)
Isotype
IFN-gama
0.47
CD8 (APC)
IFN-γ (PE)
IFN-γ PE Ab
IFN-gama
8.85
CD4 (PerCP-Cy5.5)
IFN-γ (PE)
IFN-γ PE Ab
IFN-gama
8.14
CD8 (APC)
IFN-γ (PE)
b
Lymphocytes
86.5
SSC-A :: SSC-A
FSC-A :: FSC-A
Single Cells
93.0
FSC-H :: FSC-H
FSC-A :: FSC-A
Live
99.8
SSC-H :: SSC-H
Live (eFluro780)
CD45
96.0
SSC-H :: SSC-H
CD45 (FITC)
Q1
9.06
Q2
0.16
Q4
80.5
Q3
10.3
CD4 (PerCP-Cy5.5)
CD8 (APC)
IFN-gama
8.85
CD4 (PerCP-Cy5.5)
IFN-γ (PE)
IFN-gama
8.14
CD8 (APC)
IFN-γ (PE)

**ESM Figure 8.** (**a**) Isotype control gating used in flow cytometry analysis to measure intracellular IFN-γ production in $CD4^+$ and $CD8^+$ T cells. (**b**). Flow cytometry gating strategy used to analyze intracellular IFN- γ and activation markers (CD25, CD69 and LAG-3) expression on $CD4^+$ and $CD8^+$ positive T cells. Lymphocytes were first gated on forward scatter (FSC) and side scatter (SSC), followed by gating on live $CD45^+CD4^+$ and $CD45^+CD8^+$ T cells to assess the expression intracellular IFN- γ cytokine and activation marker (CD25, CD69 and LAG-3) express cells population.
